# Mucus polymer concentration and *in vivo* adaptation converge to define the antibiotic response of *Pseudomonas aeruginosa* during chronic lung infection

**DOI:** 10.1101/2023.12.20.572620

**Authors:** Matthew A. Greenwald, Suzanne L. Meinig, Lucas M. Plott, Cristian Roca, Matthew G. Higgs, Nicholas P. Vitko, Matthew R. Markovetz, Kaitlyn R. Rouillard, Jerome Carpenter, Mehmet Kesimer, David B. Hill, Jonathan C. Schisler, Matthew C. Wolfgang

**Affiliations:** Department of Microbiology and Immunology, University of North Carolina, Chapel Hill, North Carolina, USA; Marsico Lung Institute, University of North Carolina, Chapel Hill, North Carolina, USA; Joint Department of Biomedical Engineering, University of North Carolina, Chapel Hill, North Carolina, USA; Department of Pharmacology, The University of North Carolina, Chapel Hill, North Carolina, USA; McAllister Heart Institute, University of North Carolina, Chapel Hill, North Carolina, USA

**Author notes:** Address correspondence to Matthew C. Wolfgang,. S.L.M., Pharmaceutical Product Development (PPD; Thermo Fisher Scientific), Morrisville, North Carolina, USA; N.P.V., Pfizer, Sanford, North Carolina, USA.

**Keywords:** *P. aeruginosa*, tobramycin, antibiotic efficacy, sputum, muco-obstructive airway disease

## Abstract

The airway milieu of individuals with muco-obstructive airway diseases (MADs) is defined by the accumulation of dehydrated mucus due to hyperabsorption of airway surface liquid and defective mucociliary clearance. Pathological mucus becomes progressively more viscous with age and disease severity due to the concentration and overproduction of mucin and accumulation of host-derived extracellular DNA (eDNA). Respiratory mucus of MADs provides a niche for recurrent and persistent colonization by respiratory pathogens, including *Pseudomonas aeruginosa*, which is responsible for the majority of morbidity and mortality in MADs. Despite high concentration inhaled antibiotic therapies and the absence of antibiotic resistance, antipseudomonal treatment failure in MADs remains a significant clinical challenge. Understanding the drivers of antibiotic recalcitrance is essential for developing more effective treatments that eradicate persistent infections. The complex and dynamic environment of diseased airways makes it difficult to model antibiotic efficacy *in vitro*. We aimed to understand how mucin and eDNA concentrations, the two dominant polymers in respiratory mucus, alter the antibiotic tolerance of *P. aeruginosa*. Our results demonstrate that polymer concentration and molecular weight affect *P. aeruginosa* survival post antibiotic challenge. Polymer-driven antibiotic tolerance was not explicitly associated with reduced antibiotic diffusion. Lastly, we established a robust and standardized *in vitro* model for recapitulating the *ex vivo* antibiotic tolerance of *P. aeruginosa* observed in expectorated sputum across age, underlying MAD etiology, and disease severity, which revealed the inherent variability in intrinsic antibiotic tolerance of host-evolved *P. aeruginosa* populations.

**Importance:** Antibiotic treatment failure in *Pseudomonas aeruginosa* chronic lung infections is associated with increased morbidity and mortality, illustrating the clinical challenge of bacterial infection control. Understanding the underlying infection environment, as well as the host and bacterial factors driving antibiotic tolerance and the ability to accurately recapitulate these factors *in vitro*, is crucial for improving antibiotic treatment outcomes. Here, we demonstrate that increasing concentration and molecular weight of mucin and host eDNA drive increased antibiotic tolerance to tobramycin. Through systematic testing and modeling, we identified a biologically relevant *in vitro* condition that recapitulates antibiotic tolerance observed in *ex vivo* treated sputum. Ultimately, this study revealed a dominant effect of *in vivo* evolved bacterial populations in defining inter-subject *ex vivo* antibiotic tolerance and establishes a robust and translatable *in vitro* model for therapeutic development.

## Introduction

Muco-obstructive airway diseases (MADs) comprise a spectrum of hereditary and acquired disorders that predispose individuals to recurrent and chronic bacterial lung infections. MADs are characterized by the accumulation of dehydrated airway mucus, in diseases including cystic fibrosis (CF), non-CF bronchiectasis (NCFB), primary ciliary dyskinesia (PCD), and chronic obstructive pulmonary disease (COPD) (1). Healthy airway mucus comprises a thin, well-hydrated film lining the luminal surface of the airway epithelium, which is continuously cleared by ciliary beat and cough (2–4). In MADs, defective ion–fluid transport, excessive mucin secretion, reduced mucociliary clearance, and inflammation lead to hyperconcentrated mucus accumulation and obstruction in the airways (1, 4–7). This non-functional (pathological) mucus in MADs provides a niche for colonization by commensal and pathogenic bacteria (8–13). Recurrent or chronic infection with the opportunistic Gram-negative bacterium *Pseudomonas aeruginosa* is common and becomes dominant in late-stage disease (11, 13–15). *P. aeruginosa* colonizes and persists within pathological respiratory mucus despite a robust inflammatory response and frequent antibiotic treatment (16–18). A clinical breakthrough in MAD treatment was the development of aerosolized antibiotics, allowing high-dose delivery directly to the site of infection with reduced systemic toxicity (19–21). Tobramycin, an aminoglycoside antibiotic, is one of the most commonly prescribed treatments for respiratory infections caused by *P. aeruginosa* (22–24). Inhaled tobramycin achieves an average C_max_ of 300 µg/mL in sputum; in some reports, higher concentrations have been observed (21, 25). While the introduction of inhaled antibiotic therapies has improved the quality of life and extended the life expectancy of people with MADs, antibiotic treatment failure remains a significant barrier to the resolution of infection (26). Despite the global threat of antimicrobial resistance, antibiotic treatment failure in MADs frequently occurs in the absence of observed *in vitro* antimicrobial resistance (27–33). Rather, antibiotic tolerance and persistence, bacterial physiological states that reduce the sensitivity of a population to antimicrobial killing, account for treatment failures in MADs (34–36). Moreover, antibiotic tolerance is associated with the subsequent emergence of antibiotic resistance (37, 38). Understanding the mechanisms underlying antibiotic tolerance is central to the development of effective antipseudomonal therapies for MADs.

Mucus associated with MADs is dominated by two biopolymers: mucins and host-derived extracellular DNA (eDNA). Mucins are high molecular weight (HMW; 0.5 – 20 MDa monomers) glycoproteins densely decorated with O-linked glycans and cross-linked by disulfide bonds, creating an entangled gel-like mesh network, thereby altering the biophysical properties of the mucus (5, 39, 40). eDNA in mucus primarily consists of large fragments (> 10 kb; 6 MDa) of host genomic DNA, which accumulate in the airway due to pervasive and non-productive inflammation associated with MADs (41, 42). Increased concentrations of both eDNA and mucin have been associated with subject age, clinical exacerbation, and declining lung function (11, 42, 43). Likewise, age, acute exacerbation, and disease severity predispose individuals to higher instances of antibiotic treatment failure and chronic infections (11, 34, 44–48). Given the apparent association between mucus polymer concentration, disease severity, and treatment failure, we investigated the effects of mucin and eDNA concentrations on antibiotic recalcitrance of *P. aeruginosa* using a disease-relevant *in vitro* model system.

Studying pathogen behavior in MADs has proven challenging due to the nutritional and rheological complexity of the *in vivo* environment. Laboratory media, such as lysogeny broth (LB) and Mueller-Hinton broth (MHB), often used for *in vitro* antibiotic testing, fail to recapitulate the complex *in vivo* infection microenvironment (49–54). Previous studies investigating the effect of airway mucus polymers on bacterial behaviors have traditionally relied on: laboratory media, focused on the presence or absence of mucus polymers (55), and/or utilized non-biological polymers, such as polyethylene glycol (56) or agar gels (57). Using synthetic CF sputum medium (SCFM2), a well-characterized and chemically defined *in vitro* CF mucus simulant (49–51, 53), as a starting point, we investigated the contribution of increasing concentrations of mucin and eDNA to antipseudomonal antibiotic efficacy.

We provide evidence of a concentration-dependent relationship between mucus polymers in the airway milieu and *P. aeruginosa* antibiotic survival. Furthermore, we demonstrated that mucolytic therapeutics that reduce or degrade HMW mucin and eDNA polymers, respectively, have the potential to mitigate polymer-driven tobramycin tolerance. Applying our understanding of polymer concentration-dependent effects on *P. aeruginosa* antibiotic tolerance, we assessed the utility of *in vitro* models to recapitulate antibiotic efficacy in *ex vivo* treated sputum. We identified an *in vitro* condition capable of recapitulating the antibiotic tolerance of clinical *P. aeruginosa* populations compared to *ex vivo* treated MADs sputum in a cross-section of individuals with MADs. Lastly, by using a single, disease matched model system, we demonstrate that antibiotic tolerance varies significantly across *P. aeruginosa* populations that evolve naturally within the airways.

## Results

### Antibiotic reduction of *P. aeruginosa* burden in *ex vivo* sputum is highly variable between subjects and independent of genetic resistance

As a surrogate for clinical treatment efficacy, we assessed *P. aeruginosa* survival in *ex vivo* MAD sputum following 24 h treatment with vehicle control (PBS) or high-dose tobramycin (300 μg/mL) to mimic inhaled antipseudomonal antibiotic concentrations typically achieved in the lungs (21). Fresh, spontaneously expectorated sputum was collected from 23 subjects with CF, NCFB, or PCD and a prior history of *P. aeruginosa* infection during outpatient visits (**Table 1**). Tobramycin treatment resulted in a variable reduction in the *P. aeruginosa* burden across subject samples (**Fig. 1A; Fig. S1**), capturing the heterogeneity in treatment response observed clinically (16–18). Samples from two subjects (9%) exhibited bacterial clearance below the limit of detection (indicated by #). In contrast, sixteen samples (70%) showed less than a 3-log (99.9%) reduction in bacterial burden, a common threshold for evaluating effective antimicrobial clearance (59). No correlation was observed between *P. aeruginosa* burden reduction and clinical score (percent predicted forced expiratory volume in 1 s (ppFEV_1_)) or demographic characteristics (age, sex, and race).

**Figure 1.**
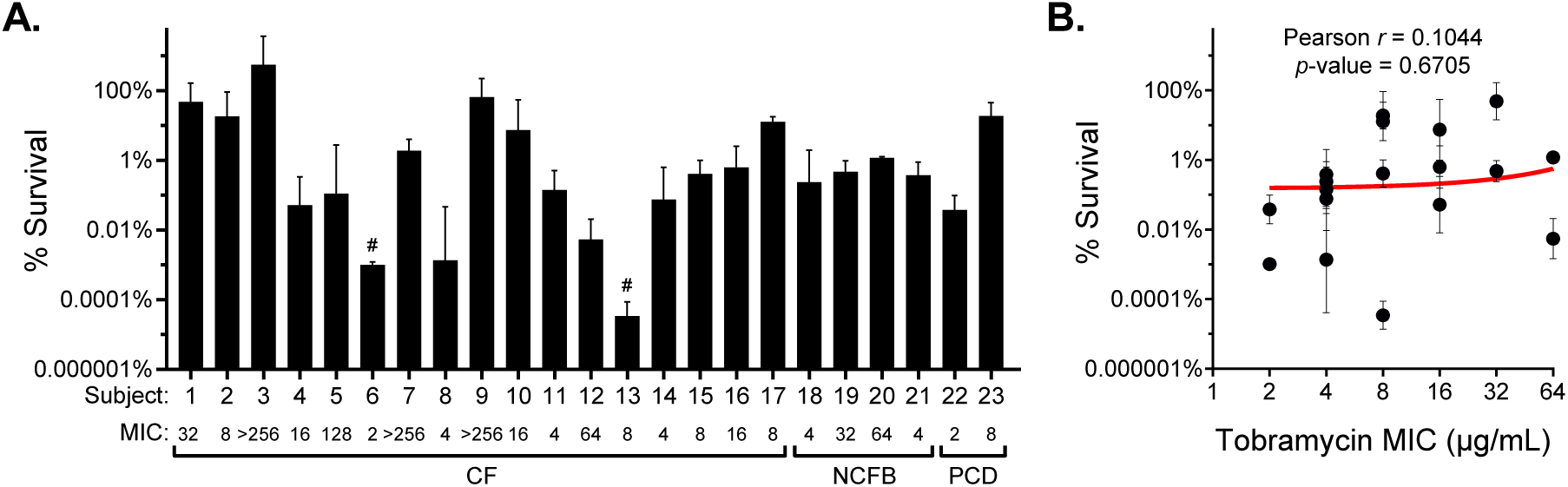
*Ex vivo* tobramycin efficacy is highly variable and independent of antibiotic resistance. **(A)** Percent survival of *P. aeruginosa* in fresh spontaneous expectorated sputum from individuals with CF, NCFB, or PCD after *ex vivo* treatment with 300 µg/mL tobramycin for 24 h. Data are presented as the percent of CFUs recovered on *Pseudomonas* isolation agar (PIA) post tobramycin treatment compared to vehicle (PBS) control. Values are reported as mean ± SD of ≥ 2 aliquots of each sputum sample tested. # indicates no CFUs recovered after tobramycin treatment (limit of detection, 4 x 10^1^ CFU/mL). The tobramycin MIC (µg/mL) for each untreated *P. aeruginosa* population is indicated. **(B)** Linear regression and Pearson correlation of *P. aeruginosa* population survival in *ex vivo* treated sputum versus population MIC, excluding hyperresistant populations (MIC ≥ 80).

**Table 1.** Demographic characteristics of sputum-producing participants.

| **Parameter** | **All**<br>*n* = 23 | **CF**<br>*n* = 17 (73.9%) | **NCFB**<br>*n* = 4 (17.4%) | **PCD**<br>*n* = 2 (8.7%) |
| --- | --- | --- | --- | --- |
| **Age in years**, med (IQR) | 37 (23.5 - 46) | 25 (22 - 37) | 70 (66.8 - 72) | 50 (49.5 - 50.5) |
| **Female**, *n* (%) | 13 (56.5%) | 9 (52.9%) | 2 (50%) | 2 (100%) |
| **Race**, *n* (%) |  |  |  |  |
| Caucasian/White | 16 (69.6%) | 12 (70.6%) | 3 (75%) | 1 (50%) |
| African American/Black | 7 (30.4%) | 5 (29.4%) | 1 (25%) | 1 (50%) |
| **CFTR<sup>a</sup> genotype**, *n* (%) |  |  |  |  |
| F508del homozygous | - | 7 (41.2%) | - | - |
| F508del heterozygous | - | 6 (35.3%) | - | - |
| No F508del mutation | - | 4 (23.5%) | - | - |
| **ppFEV<sub>1</sub><sup>b</sup>**, mean ± SD | 51.9 ± 15.7 | 53.6 ± 14.3 | 54.3 ± 19.3 | 34.3 ± 4.5 |

<sup>a</sup>CFTR; cystic fibrosis transmembrane conductance regulator
<sup>b</sup>ppFEV<sub>1</sub>; percent predicted forced expiratory volume in 1 s

To determine whether antibiotic resistance was associated with bacterial burden reduction, tobramycin susceptibility was measured by Etest using whole *P. aeruginosa* populations recovered from mock-treated sputum. Sixteen samples (70%) harbored *P. aeruginosa* with clinically defined resistance (MIC ≥ 8 μg/mL) (60); of these, four harbored *P. aeruginosa* hyperresistant to tobramycin (MIC ≥ 80 μg/mL, 10x the clinical resistance breakpoint (60)). Excluding subjects with hyper-resistant *P. aeruginosa* (*n* = 4), no correlation (*r* = 0.1044; *p*-value = 0.6705) was observed between *ex vivo* tobramycin survival and *in vitro* population MIC (**Fig. 1B**), which is consistent with previous reports of a disconnect between resistance and bacterial clearance in MADs (22, 61, 62).

### *P. aeruginosa* tobramycin tolerance increases with mucus polymer concentration *in vitro*

In people with MADs, the concentration of mucin and eDNA in airway mucus increases with age, disease severity, and during acute exacerbation (11, 42, 43). Similarly, age, exacerbation status, and disease progression are associated with antibiotic treatment failure and persistent pathogen colonization (11, 34, 44–48). To explore the impact of increasing mucin and eDNA polymer concentrations on *P. aeruginosa* behavior, we used increasing concentrations of partially purified Muc5AC porcine gastric mucin (mucin) and HMW salmon sperm DNA (eDNA) in SCFM2. The growth kinetics of the laboratory strain mPAO1 were not affected by the addition of either polymer across the biologically relevant range tested (**Fig. S2**). To determine how mucus polymer concentration influences *P. aeruginosa* survival following antimicrobial treatment, we grew mPAO1 for 8 h, to early stationary growth phase, in SCFM2 with and without increasing concentrations of mucin and eDNA, followed by challenge with tobramycin (300 μg/mL) for 24 h (**Fig. 2**). *P. aeruginosa* was completely eradicated in lysogeny broth (LB) and SCFM2 lacking mucin and eDNA. At healthy/mild disease mucin concentrations (0.5% and 1% w/v mucin) (11), *P. aeruginosa* was eradicated or reduced by > 6-logs, respectively. Increasing mucin concentration to 2% resulted in a 3-log increase in bacterial survival, relative to 1% (**Fig. 2A**). Notably, mucin concentrations ≥ 2% are consistent with mucus hyperconcentration observed in muco-obstructive disease (11). Survival further increased at 3% and 4% mucin, and a maximum bacterial survival plateau was reached at ≥ 4% mucin where only 1-log of killing was observed (**Fig. 2A**). Mucin concentration-dependent antibiotic tolerance was similarly observed for ciprofloxacin (300 μg/mL) (**Fig. S3A**). (63). The emergence of antibiotic resistance was not observed in any experiment, as evaluated by the agar dilution method (75).

**Figure 2.**
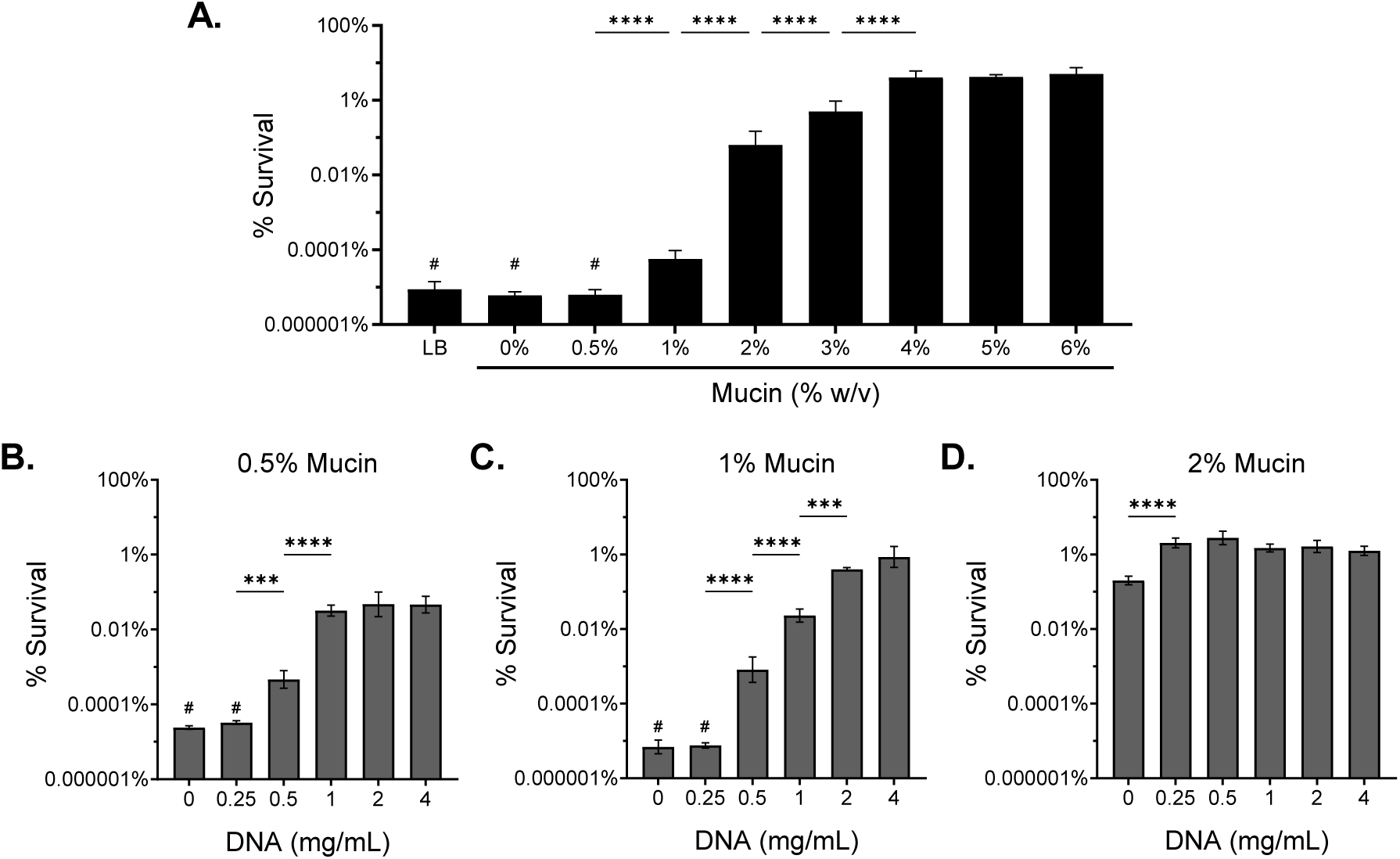
Increasing mucus polymer concentration drives antibiotic tolerance to tobramycin. Percent survival of mPAO1 after 24 h treatment with 300 µg/mL tobramycin in **(A)** LB and SCFM2 with increasing % w/v mucin without eDNA or SCFM2 at a consistent mucin concentration of **(B)** 0.5%, **(C)** 1%, or **(D)** 2% w/v with increasing HMW eDNA. Data represent the mean ± SD of *n* ≥ 3 biological replicates. Statistical differences were analyzed using one-way ANOVA with Tukey’s multiple comparison correction. \*\*\**p*<0.001, \*\*\*\**p*<0.0001. # indicates no *P. aeruginosa* recovered after antibiotic treatment (limit of detection, 10^2^ CFU/mL).

Next, we evaluated a range of HMW eDNA concentrations from 0 to 4 mg/mL, representing healthy to severe disease concentrations reported for people with CF (64), in SCFM2 at multiple mucin concentrations. Increasing eDNA in SCFM2 containing 0.5% and 1% mucin resulted in a concentration-dependent increase in *P. aeruginosa* survival following antibiotic treatment (**Fig. 2B and C**). However, the effect of increasing eDNA concentration on bacterial survival was only observed at low mucin concentrations. The addition of eDNA above 0.25 mg/mL in 2% mucin SCFM2 resulted in no further increase in bacterial survival (**Fig. 2D**). Furthermore, eDNA polymer size was crucial for driving tolerance, as low molecular weight (LMW) eDNA (< 200 bp) (**Fig. S4A**) did not increase tolerance, regardless of concentration (**Fig. S4B and C**). Overall, we observed that the concentrations of both mucin and HMW eDNA significantly affected the efficacy of tobramycin treatment against *P. aeruginosa in vitro*.

### Reduction of mucus polymer size improves antibiotic efficacy

Based on the findings that mucus polymer concentration (mucin and eDNA) and size (eDNA) affect antibiotic killing of *P. aeruginosa*, we assessed the utility of mucolytics, which function to reduce or degrade mucus polymers, to improve tobramycin efficacy. Recombinant human DNase I (rhDNase I; Pulmozyme, Dornase Alfa) is a commonly prescribed mucolytic that degrades eDNA and improves mucus clearance from the lungs of individuals with MADs (41, 65, 66).

The addition of rhDNase to SCFM2 at the average sputum C_max_ after administration (3 µg/mL) (67) degraded 1 mg/mL of HMW eDNA in 2 h at 37°C (**Fig. S4D**). Additionally, rhDNase alone did not have direct antimicrobial activity or alter *P. aeruginosa* burden (**Fig. S4E**). To assess the effect of rhDNase on antibiotic efficacy, rhDNase was added at various time points after *P. aeruginosa* inoculation in SCFM2 containing 0.5% or 2% mucin and 1 mg/mL HMW eDNA and challenged with tobramycin. We observed that the addition of rhDNase at the time of bacterial inoculation (0 h) and up to 4 h after bacterial inoculation resulted in a significant > 3-log reduction in *P. aeruginosa* survival in SCFM2 containing 0.5% mucin (**Fig. 3A**), and a significant 1-log reduction in 2% mucin (**Fig. 3B**) 24 h post-antibiotic challenge compared to the no rhDNase control (dark gray bar). The resulting level of bacterial survival when rhDNase was added early (≤ 4 h) was similar to the level of tolerance observed in SCFM2 lacking eDNA. The effect of rhDNase on decreasing tobramycin survival was intermediate or negligible when added at 6 or 8 h, respectively.

**Figure 3.**
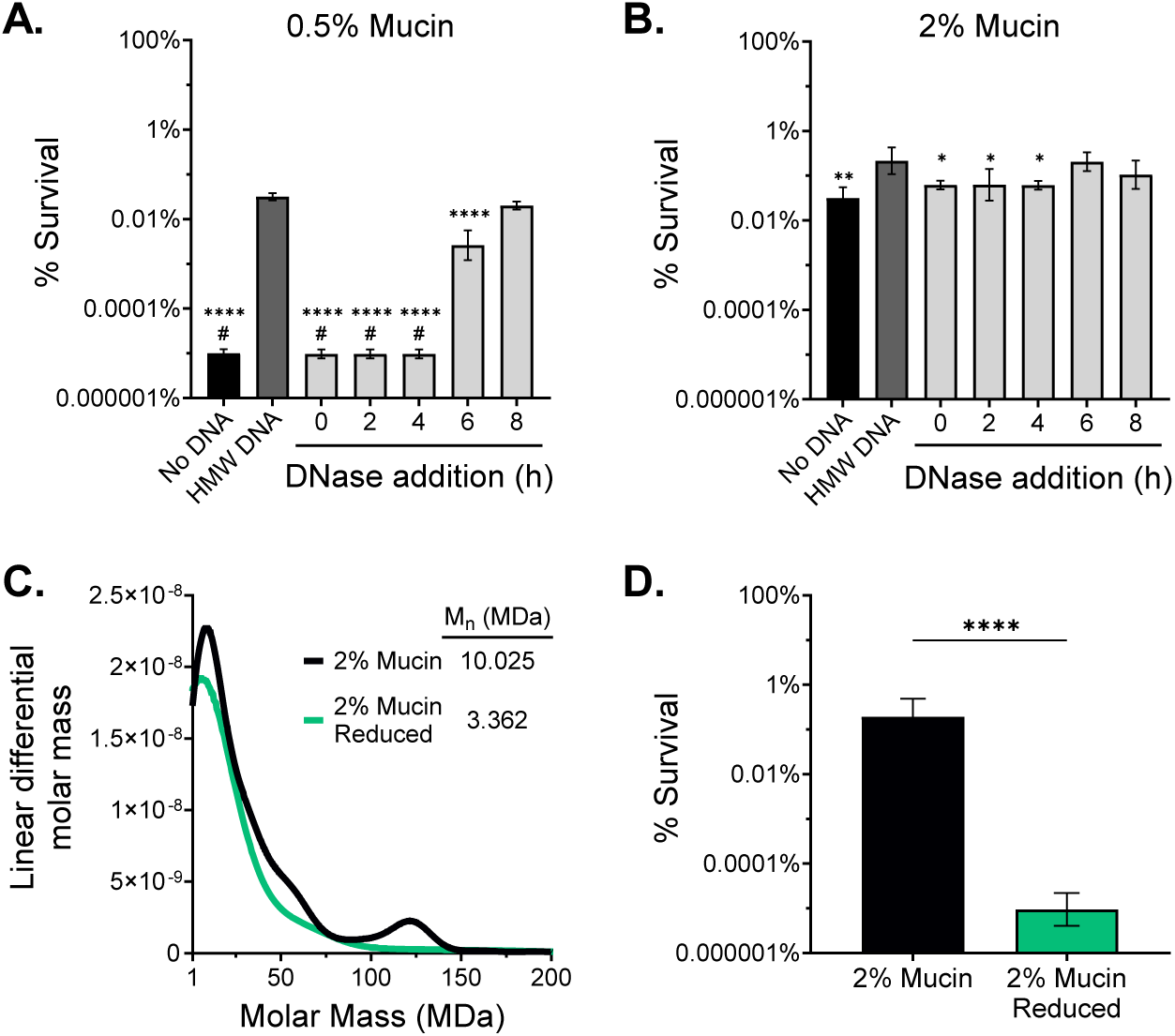
Polymer size reduction improves tobramycin efficacy. Survival of mPAO1 after tobramycin challenge in **(A)** 0.5% or **(B)** 2% w/v mucin SCFM2 with no eDNA, 1 mg/mL HMW eDNA, or 1 mg/mL HMW eDNA with rhDNase (3 μg/mL) added at various time points after bacterial inoculation (0 h). Significant differences were determined by one-way ANOVA with Dunnett’s correction relative to the HMW eDNA control (dark gray bar). \**p*<0.05, \*\**p*<0.01, \*\*\*\**p*<0.0001. #, no viable CFU after antibiotic treatment (limit of detection, 10^2^ CFU/mL). **(C)** Molar mass distribution from SEC-MALS of dialyzed 2% w/v mucin before and after treatment with 25 mM DTT, 4 M guanidine, and 50 mM iodoacetamide. M_n_; number average molecular weight. **(D)** mPAO1 survival to tobramycin in 2% reduced mucin SCFM2. Statistical significance was evaluated by a two-tailed Student’s *t*-test. \*\*\*\**p*<0.0001. Data presented as mean ± SD of *n* ≥ 3 biological replicates.

While eDNA targeted therapies have the potential to improve antibiotic efficacy in MADs, our data suggest that at disease-relevant concentrations (≥ 2%) antibiotic tolerance is dominated by mucin. In contrast to eDNA, mucins present a greater therapeutic challenge. Mucin monomers form a mesh-like network resulting from the formation of intermolecular disulfide bonds, which greatly increases the effective molecular weight of the polymer and alters its biophysical properties (5). Disruption of disulfide cross-linked mucins by reducing agents improves mucociliary clearance and airway obstruction (22). To determine whether porcine gastric mucin (PGM) contained multimers that could be reduced, we assessed polymer molecular weight using size-exclusion chromatography coupled with multi-angle laser light scattering (SEC-MALLS) before or after treatment with dithiothreitol (DTT), a potent reducing agent, combined with guanidine to improve *in vitro* reducing activity, and iodoacetamide to prevent disulfide bond reformation. The partially purified mucin was confirmed to contain multimers that were eliminated upon mucolytic reduction, as evidenced by the decrease in the number average molecular weight fraction from 10 to 3 MDa, and by the elimination of a HMW (120 MDa) multimer peak between unreduced (black) and fully reduced (green) mucin (**Fig. 3C**). The antibiotic tolerance of *P. aeruginosa* in SCFM2 containing reduced 2% mucin was significantly decreased by > 4-log compared to SCFM2 with an equivalent concentration of unreduced mucin (**Fig. 3D**). Together, these data demonstrate the importance of both polymer concentration (**Fig. 2**) and molecular weight (**Fig. 3; Fig. S4**) in driving antibiotic tolerance of *P. aeruginosa* in the MAD mucus environment.

### The effect of mucus polymers on antibiotic diffusion and sequestration does not explain *P. aeruginosa* tolerance to tobramycin

To evaluate the possibility that restricted diffusion from increased mucin and eDNA concentrations accounted for the reduced antimicrobial activity of tobramycin, we measured the diffusion of tobramycin through SCFM2 as a function of polymer concentration. We observed no significant effect of increasing concentrations of either mucin or eDNA on the diffusion rate of Texas Red conjugated tobramycin (**Fig. S5**). These data suggested that reduced tobramycin diffusion was not responsible for the observed changes in antibiotic tolerance.

To rule out tobramycin sequestration by mucin or eDNA binding, we performed broth microdilution MIC in a gradient of mucin and eDNA concentrations. Similar to previous studies, we observed a mucin-dependent effect of tobramycin on the MIC against *P. aeruginosa* (68). Mucin at 0.5% and 1% w/v increased the MIC 4-fold relative to that of MHB alone. Higher mucin concentrations (2% and 4%) further increased the MIC to 8 μg/mL and 32 μg/mL, respectively (**Table 2**). At all concentrations tested, eDNA did not alter the tobramycin MIC (**Table 2**). These data suggest that mucin, but not eDNA, may partially contribute to antibiotic treatment failure through the sequestration of tobramycin. However, reduced mucin retained the same antibiotic sequestration properties as unreduced mucin (**Table 2**), even though reduced mucin conferred significantly less tobramycin tolerance than unreduced mucin (**Fig. 3D**). Despite an increase in the overall tobramycin MIC with increasing mucin concentration, the MIC at the highest mucin concentration (4%) remained approximately 10-fold lower than the concentration of tobramycin used in killing experiments. In contrast, polymyxin B and colistin, positively charged cationic antimicrobial peptides known to bind negatively charged mucin and eDNA polymers (69), exhibited significantly reduced antibiotic activity when assessed by both MIC and antibiotic survival assays at all mucin concentrations tested (**Table 2; Fig. S3B and C**).

**Table 2.** *In vitro* broth microdilution MIC in the presence of mucin or DNA*^a^*.

| Antibiotic | *In vitro* conc.<sup>b</sup> | Resistance Breakpoint<sup>c</sup> | MIC Media Condition |  |  |  |  |  |
| --- | --- | --- | --- | --- | --- | --- | --- | --- |
|  |  |  | 0% Mucin | 0.5% Mucin | 1% Mucin | 2% Mucin | 3% Mucin | 4% Mucin |
| Tobramycin | 300 | ≥ 8 | 1 | 4 | 4 | 8 | 16 | 32 |
| Ciprofloxacin | 300 | ≥ 2 | 0.5 | 0.5 | 0.5 | 0.5 | 0.5 | 0.5 |
| Colistin | 40 | ≥ 4 | 2 | 32 | 64 | 128 | 256 | 256 |
| Polymyxin B | 40 | ≥ 4 | 2 | 64 | 128 | 256 | 256 | 256 |
|  |  |  | 0 mg/mL DNA | 0.25 mg/mL DNA | 0.5 mg/mL DNA | 1 mg/mL DNA | 2 mg/mL DNA | 4 mg/mL DNA |
| Tobramycin | 300 | ≥ 8 | 1 | 1 | 1 | 1 | 1 | 1 |
| Ciprofloxacin | 300 | ≥ 2 | 0.5 | 0.5 | 0.5 | 0.5 | 0.5 | 0.5 |
| Colistin | 40 | ≥ 4 | 2 | 2 | 2 | 4 | 4 | 16 |
| Polymyxin B | 40 | ≥ 4 | 2 | 2 | 2 | 4 | 16 | 32 |
|  |  |  | 0% Mucin Reduced | 0.5% Mucin Reduced | 1% Mucin Reduced | 2% Mucin Reduced |  |  |
| Tobramycin | 300 | ≥ 8 | 1 | 4 | 8 | 16 |  |  |

<sup>a</sup>All antibiotic concentrations and MIC values are in µg/mL
<sup>b</sup>Antibiotic concentration used in this study
<sup>c</sup>CLSI breakpoints (60)

### Mucus polymer concentration drives viscous rheology

Previous studies identified mucus rheology as a driver of antibiotic treatment failure (46, 47, 70–73). To determine whether the increased polymer concentration influenced the viscoelasticity of the media and antibiotic efficacy, we analyzed the biophysical properties of SCFM2 across a mucin and eDNA concentration gradient. Complex viscosity (*η*\*) measures the ability of a particle to pass through a medium and the resistance of that medium to deformation. In sputum, high viscosity can reduce the diffusion and movement of therapeutics and bacteria, in addition to the sequestration of molecules within the mucus network (27, 68, 74). As expected, the measured viscoelasticity of the medium increased with the mucin content (**Fig. 4A)**, and HMW eDNA concentration (**Fig. 4B and C**). LMW DNA contributed no viscoelasticity to the medium (**Fig. S4F and G**). Reduction of mucin multimers in the mucin preparations by DTT treatment significantly reduced the viscoelasticity of the sample when assayed at matching concentration (**Fig. 4D**). These results are consistent with those of previous studies on both in-tact *ex vivo* mucus and purified commercial mucins, showing concentration-dependent rheological properties (5, 7, 45–47, 75).

**Figure 4.**
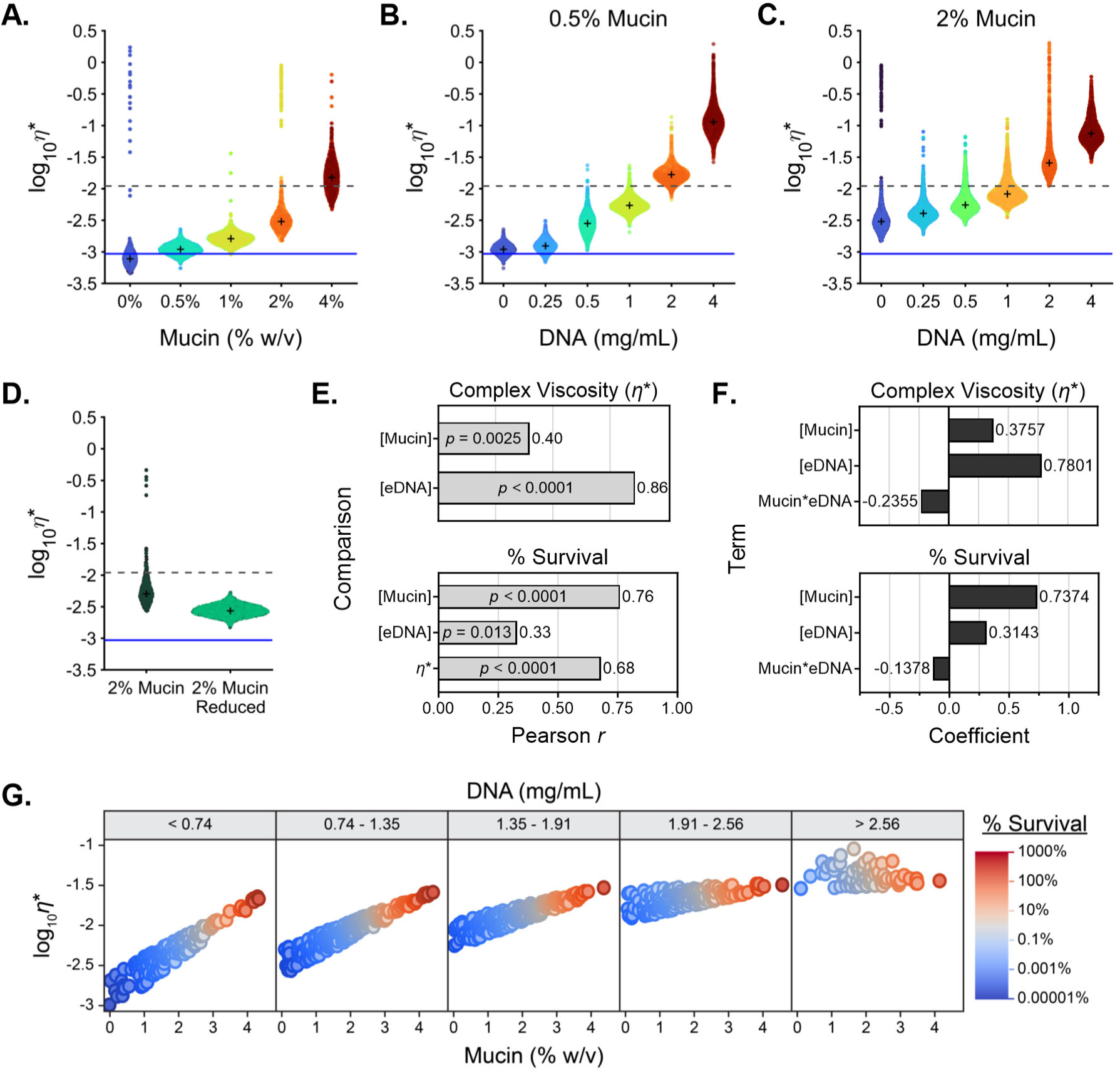
Biopolymer concentration drives viscoelasticity but does not fully explain tobramycin tolerance. **(A-D)** Modulus of the complex viscosity (*η*\*) of a 1 µm diameter microsphere moving through **(A)** LB and SCFM2 with increasing concentrations of mucin (% w/v) without eDNA, increasing eDNA concentrations (mg/mL) in **(B)** 0.5% and **(C)** 2% w/v mucin SCFM2, or **(D)** 2% mucin SCFM2 before and after mucolytic reduction. Data are representative of *n* ≥ 3 technical replicates. The viscosity of water is indicated by the solid blue line and the complex viscosity at which mucus entangles and becomes gel-like (40) is indicated by the dashed gray line. + indicates the median for each sample. **(E)** Multivariate correlation of viscoelasticity to mucin and DNA concentration, and survival to mucin, DNA concentration, and viscoelasticity, represented by bar graph of Pearson’s *r* with the corresponding *p*-value within each bar. **(F)** Centered and scaled coefficients for the X variables used for partial least squares regression in modeling viscoelasticity and survival to tobramycin. **(G)** Visualization of the simulation results via grouped dot plots.

### Mucin concentration is the dominant driver of antibiotic tolerance

Our data indicate that mucins and eDNA affect antibiotic tolerance of *P. aeruginosa* and the viscoelasticity of the medium. We used a multivariant analysis to measure the linear relationships across and between the outcome variables (tobramycin survival and viscoelasticity) and input variables (mucin and eDNA concentration) (**Fig. 4E; Fig. S6 A-F**). While both mucin and eDNA concentration showed a positive correlation to viscoelasticity, eDNA concentration showed the strongest correlation (*r* = 0.86; *p*-value = 8.76E-17), suggesting eDNA is the primary driver of viscoelasticity. Similarly, [mucin], [eDNA], and viscoelasticity each demonstrate a positive correlation with antibiotic survival, yet the highest correlation (*r* = 0.76; *p*-value = 2.34E-11) was observed with mucin concentration (**Fig. 4E**). Given the multicollinearity between and among the outcome, or “Y” variables (tobramycin survival and viscoelasticity), and input, or “X” variables (mucin and eDNA concentration), we used partial least squares linear regression to model our experimental system (76). With a three-factor model, we can explain 86% of the variation in survival and viscoelasticity using [mucin], [eDNA], and the interaction term (mucin*eDNA) (**Fig. 4F**).

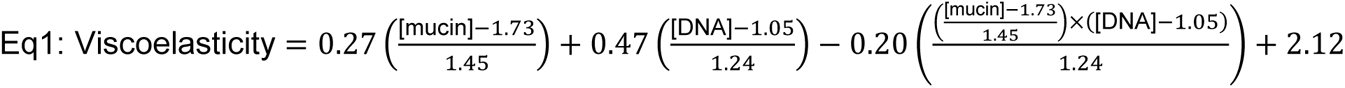

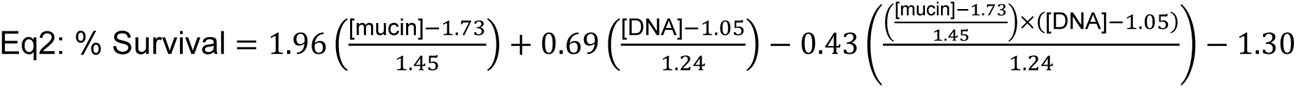

The model performed well for both outcome variables, with R^2^ values of 0.98 and 0.73 for viscoelasticity and tobramycin survival, respectively (**Fig. S6G and H**), and revealed that mucin concentration had the highest coefficient for determining tobramycin antibiotic survival, whereas eDNA concentration had the highest coefficient for driving viscoelasticity (**Fig. 4F**). The trends of the interactions between mucin and eDNA indicated that the outcome variables are differentially affected by the mucin*eDNA interaction (**Fig. 4F**). Next, we built a predictor profiler, available at https://public.jmp.com/spaces/ALLUSERS/posts/3gyw4Hl2FCmqM648mJdGm/reports, and paired that with Monte Carlo simulations using a truncated normal distribution of the experimental range of eDNA and mucin used in our study. We visualized the simulation results via grouped dot plots (**Fig. 4G**) and individual biplots (**Fig. S6 I-N**). Most notably, tobramycin survival was positively associated with mucin across all eDNA concentrations. In contrast, the positive association between mucin and viscoelasticity weakened as eDNA concentration increased. This differential effect of moderate to high levels of eDNA in the simulation was also seen with the dose-dependent decrease in the positive correlation between viscoelasticity and survival (**Fig. 4G**). Together, these models establish that mucin polymer concentration is the dominant driver of *P. aeruginosa* recalcitrance to antibiotic treatment but can be further modified by the amount of eDNA present.

### *In vitro* testing reveals a single condition that recapitulates *ex vivo* sputum antibiotic efficacy and heterogeneous *P. aeruginosa* population-level tolerance

Unlike laboratory strains, *P. aeruginosa* clinical isolates from individuals with MADs exhibit evolved behaviors from adaptation to the host mucus environment and repeated antibiotic exposure (14, 15, 77). To determine whether our observations, using a laboratory strain to model antibiotic tolerance, were recapitulated in clinical isolates, we assessed the *in vitro* tobramycin survival of total *P. aeruginosa* populations isolated from expectorated sputum. We seeded SCFM2 with recovered sputum *P. aeruginosa* populations isolated from mock-treated sputum assayed in **Fig. 1A** to assess survival post-antibiotic challenge *in vitro*. We focused on the effects of mucin concentration, based on the dominant effect of mucin on antibiotic tolerance (**Fig. 4E-G**). All experiments were performed with 1 mg/mL HMW eDNA, the average eDNA concentration associated with pathological sputum (5) (**Fig. S7**). For these studies, we excluded the four hyper-resistant populations (tobramycin MIC ≥ 80 μg/mL, **Fig. 1A**). Similar to the results obtained with mPAO1 (**Fig. 2A**), all clinical populations of *P. aeruginosa* demonstrated a form of mucin concentration-dependent tobramycin tolerance (**Fig. 5A-C; Fig. S8**); however, there was substantial variability in the baseline population tolerance, highlighting the impact of bacterial *in vivo* evolution driving differential antibiotic efficacy. These data suggest that while polymer concentration greatly contributes to antibiotic tolerance, inherent inter-population variability is the dominant driver of survival in the face of high-dose tobramycin challenge.

**Figure 5.**
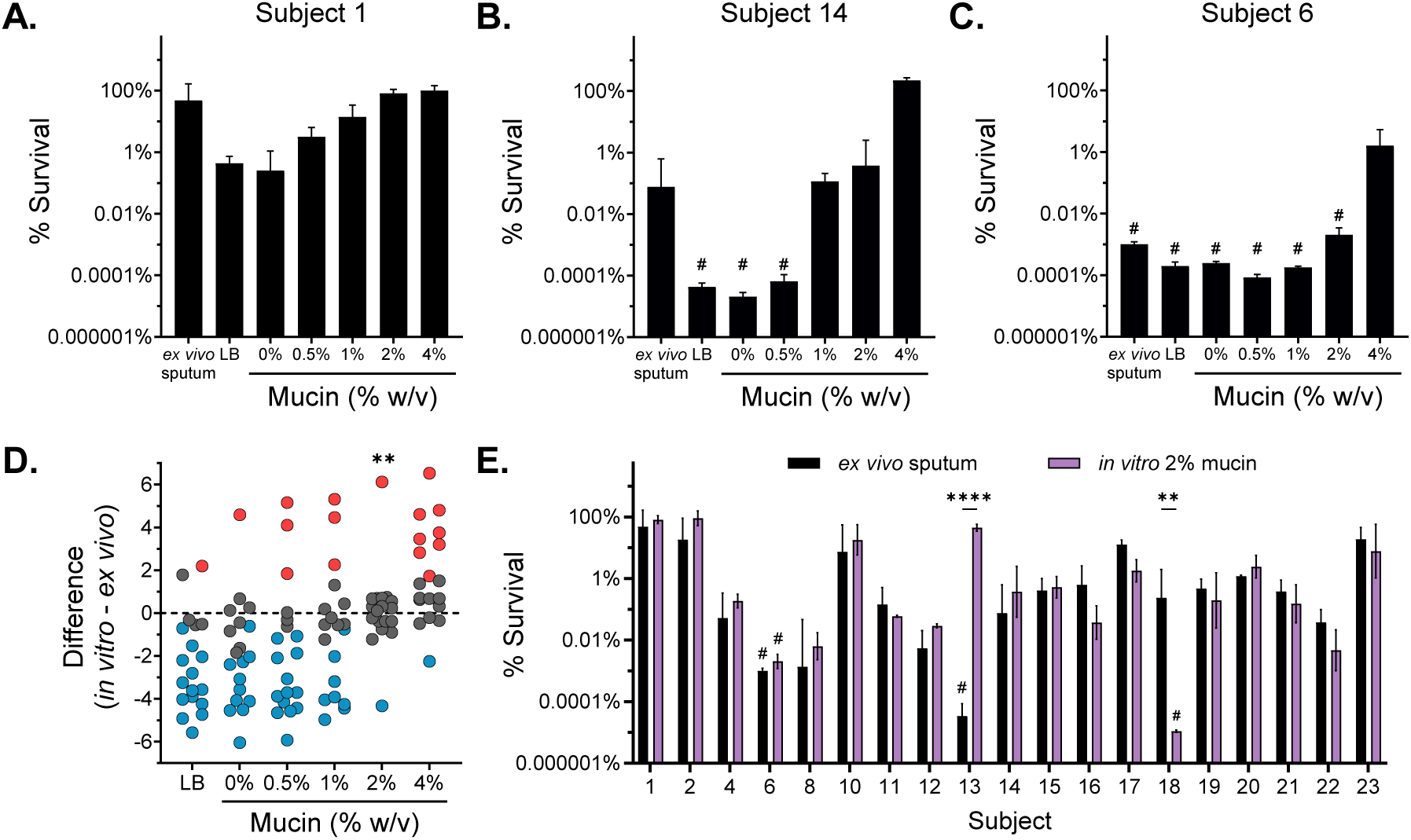
Clinical populations of *P. aeruginosa* demonstrate mucin concentration-dependent killing and reveal a relevant model of *in vivo* antibiotic efficacy. Survival of *P. aeruginosa* populations isolated from fresh expectorated sputum cultured *in vitro* in LB and SCFM2 at 1 mg/mL eDNA with increasing concentrations of mucin and treated for 24 h with 300 µg/mL tobramycin, excluding hyperresistant populations. (A-C) *P. aeruginosa* population survival of a representative **(A)** high-, **(B)** medium-, and **(C)** low-tolerant sputum populations all demonstrate mucin concentration-dependent antibiotic tolerance *in vitro*. See **Fig. S8** in the supplementary material for the entire dataset. **(D)** Differences between the observed model *ex vivo* sputum tolerance and the experimental *in vitro* tolerance of *P. aeruginosa* populations evaluated in LB and SCFM2 with increasing % w/v mucin were evaluated by one-way ANOVA with Dunnett’s post-test (|d| = 2.9, α = 0.05). Points are colored if they overestimated (red) or underestimated (blue) *ex vivo* tolerance (*p*-value < 0.05). Analysis of means for proportions determined the *in vitro* culture conditions with a disproportionate similarity to *ex vivo* tolerance compared to the null hypothesis. \*\**p*<0.01. **(E)** Comparison of *P. aeruginosa* population survival treated *ex vivo* in sputum (black bars) and the same *P. aeruginosa* populations treated *in vitro* after growth in 2% w/v mucin SCFM2 with 1 mg/mL eDNA (purple bars). Statistical differences were determined using individual unpaired two-tailed Student’s *t*-test. \*\**p*<0.01, \*\*\*\**p*<0.0001. Data are representative of *n* = 3 independent replicates and plotted as mean ± SD. # indicates no recovered *P. aeruginosa* after antibiotic treatment (limit of detection, 10^2^ CFU/mL).

Based on our investigation of *in vitro* mucin concentration-dependent antibiotic tolerance and *ex vivo* testing, we sought to understand the predictive value of *in vitro* testing. To evaluate the power of *in vitro* conditions to recapitulate *ex vivo* antibiotic tolerance, we assessed the difference in antibiotic survival of each sputum-derived *P. aeruginosa* population grown *in vitro* in LB and SCFM2 across a range of mucin concentrations to its corresponding observed *ex vivo* treated sputum tolerance (**Fig. 5D; Fig. S9A**). A difference near zero indicates that an *in vitro* condition imparted tobramycin tolerance similar to that observed in the corresponding *ex vivo* sputum. LB and SCFM2 with mucin concentrations ≤ 1% underestimated (blue symbols) antibiotic survival compared to the observed *ex vivo* treatment, whereas a higher mucin concentration (4%) in SCFM2 generally overestimated (red symbols) *ex vivo* tobramycin survival (**Fig. 5D**). Moreover, we found that SCFM2 containing 2% mucin with 1 mg/mL HMW eDNA was the only experimental condition that mimicked *ex vivo* tolerance across the patient-derived cultures (*p*-value < 0.01) (**Fig. 5D; Fig. S9A and B**). In fact, survival between *in vitro* 2% mucin SCFM2 and *ex vivo* sputum was indistinguishable in all but two subject samples, as confirmed by individual *t*-tests (**Fig. 5E**). Thus, we identified a robust *in vitro* condition capable of recapitulating tobramycin efficacy against evolved *P. aeruginosa* populations in sputum with 89.5% accuracy, compared to an average of 43% across all other conditions (**Fig. S9B**). By identifying a single *in vitro* condition that accurately controls for the host mucus environment across subjects, we found that inherent *P. aeruginosa* population-level tolerance is the major driver of tobramycin treatment efficacy, independent of bacterial resistance.

## Discussion

*P. aeruginosa* infections in people with MADs remain a complex clinical challenge despite the development of high-dose inhaled antimicrobial delivery and highly effective modulator therapies (HEMTs) such as Kalydeco and Trikafta/Kaftrio (78, 79). Paradoxically, antipseudomonal therapies fail to clear infections, even in the absence of observed *in vitro* antimicrobial resistance (27–33). Treatment of *ex vivo* sputum from multiple subjects with MADs resulted in extreme variability in tobramycin clearance of *P. aeruginosa*, as shown in **Fig. 1A**. The majority of subject sputum samples (70%) showed a modest reduction in *P. aeruginosa* burden, indicative of antibiotic treatment failure (< 3-log reduction) (59). In spite of measured tobramycin resistance (MIC ≥ 8 μg/mL), the *in vitro* population MIC was not predictive of reduction in *P. aeruginosa* burden (**Fig. 1B**). This was true even when considering hetero-resistance (detected by enhanced *in vitro* susceptibility testing of the total *P. aeruginosa* population cultured from expectorated sputum), which is a common limitation clinically and in other studies that typically utilize a single or a limited number of isolates for susceptibility testing (80, 81). These results mirror clinical observations and exemplify why antimicrobial susceptibility testing is not a primary factor dictating antibiotic treatment selection for *P. aeruginosa* infections in MADs (22, 27–33).

The lack of agreement between *in vitro* antibiotic resistance testing and treatment outcomes across *ex vivo* subject samples led us to explore the impact of the known variability of mucin and eDNA concentrations in respiratory mucus as drivers of *P. aeruginosa* antibiotic tolerance (11, 34, 44–48). The modulation of mucin and eDNA concentrations in a modified SCFM2 formulation allowed us to investigate the contribution of these dominant mucus polymers to antibiotic treatment failure in *P. aeruginosa*. SCFM2 has previously been shown to sufficiently recapitulate the complex nutritional environment in the average person with CF and the bulk transcriptomic profile of *P. aeruginosa* within CF sputum (49–51, 53, 82). We overcame the limitations of using commercial PGM as a mucin source through partial purification by ultracentrifugation and dialysis. Ultracentrifugation eliminated microbial contaminants, insoluble aggregated mucin polymers, and other debris, while preserving intact soluble mucins (**Fig. 3C and 4A**). Dialysis serves to equilibrate salt and metal concentrations, which have been previously reported to affect *P. aeruginosa* behavior and antimicrobial efficacy (53).

Increasing mucin concentration, across a biologically relevant range, in SCFM2 correlated with increased bacterial survival post-antibiotic challenge at clinically relevant concentrations for inhaled tobramycin (**Fig. 2A**) (21) and ciprofloxacin (**Fig. S3A**) (63). Mucin concentrations representative of healthy individuals or individuals with mild disease (0.5% and 1% mucin) demonstrated significantly lower antibiotic survival than mucin concentrations associated with disease (≥ 2% mucin) (**Fig. 2A**). Similarly, HMW eDNA induced a polymer concentration-dependent increase in antibiotic survival. However, this effect was observed primarily in SCFM2 containing 0.5% or 1% mucin (**Fig. 2B and C**), with a minimal effect observed in 2% mucin (**Fig. 2D**). At lower concentrations of mucin, the polymers have more pores that can intercalate eDNA, which may account for the dominance of eDNA in promoting antibiotic tolerance at low mucin concentrations (5). These data suggest that hyperconcentration and accumulation of mucin and eDNA within a subject result in increased tolerance of *P. aeruginosa* to antibiotics. This is consistent with clinical observations of mucin and eDNA polymer concentrations and instances of antibiotic treatment failure, both increasing as a function of age, disease severity, and clinical exacerbation (11, 34, 44–48).

As a cautionary note, commercial DNA sources are not all equal. We observed variation in DNA fragment size from commercial sources (**Fig. S4A**). The effects of eDNA polymer concentration on antibiotic tolerance were only observed with HMW eDNA, suggesting that polymer size is an important driving factor underlying antibiotic treatment failure. Our data demonstrated a significant improvement in tobramycin bacterial killing by modifying the mucus environment with mucolytic agents targeting mucin and eDNA (**Fig. 3**). However, similar to previous reports (45), the impact of rhDNase on antibiotic recalcitrance was minimal at high mucin concentrations, which indicates that rhDNase may provide the greatest benefit to antibiotic outcomes earlier disease when mucin concentration is low, or in regions of the lung where mucin concentrations have not yet reached high levels. Improved antibiotic efficacy with rhDNase was dependent on rhDNase addition prior to tobramycin challenge, which may suggest that actively growing *P. aeruginosa* or adaptation to the environment may be essential for the bacterial changes that alter susceptibility to antibiotics, such as the formation of biofilms or biofilm-like aggregates (**Fig. 3A and B**). Thus, reducing the polymer size after *P. aeruginosa* acclimatization and adaptation to the environment would not significantly impact antimicrobial tolerance. Mucolytic reduction of mucin, which does not alter the concentration of mucin monomers, but rather eliminates the multimeric mucin complexes (**Fig. 3C**) and reduces the viscoelasticity of the medium (**Fig. 4D**), significantly decreasing *P. aeruginosa* survival after antibiotic challenge (**Fig. 3D**). These data provide evidence for the utility of synergistic effects of coordinating or combining already approved mucolytic treatments with antibiotics to increase bacterial clearance in people with MADs.

Neither eDNA nor mucin presented a measurable diffusion barrier to tobramycin (**Fig. S5**); however, our data are consistent with previous reports that the charged mucin polymers can sequester charged antibiotics, thereby increasing the concentration of antibiotics required (**Table 2**). While it is likely that antibiotic sequestration partially contributes to antibiotic failure, our data suggest that this does not fully explain the polymer-driven phenomenon observed in this study. First, despite driving polymer concentration-dependent antibiotic tolerance (**Fig. 2B-D; Fig. S3A**), neither mucin nor eDNA increased the MIC of ciprofloxacin and eDNA did not contribute to tobramycin sequestration (**Table 2**). Second, disulfide bond reduced mucin demonstrated a significant decrease in antibiotic tolerance (**Fig. 3D**), despite maintaining the same *in vitro* broth MIC as unreduced mucin (**Table 2**). Thus, we concluded that while increasing mucin concentration reduces tobramycin availability, the effect does not fully account for the reduced killing observed at the inhaled tobramycin C_max_ (300 μg/mL) (21).

The effect of mucin binding or sequestration is more evident for cationic antimicrobial peptides: polymyxin B and colistin (**Table 2**). Mucin has been shown to increase the MIC of polymyxin B and colistin by > 100-fold (83). In SCFM2, mucin had an absolute effect on polymyxin B and colistin, with the lowest (0.5%, healthy) mucin concentration completely eliminating the antimicrobial activity (**Fig. S3B and C**).

The complex viscosity (*η*\*) of respiratory mucus plays a crucial role in determining the behavior of bacteria within the mucus layer. The viscoelastic properties of mucus influence bacterial motility, adhesion, biofilm formation, chemotactic responses, nutrient availability, interactions with the immune system, and the response to antibiotics (5). Understanding these interactions is important to address respiratory infections and develop targeted treatments. In SCFM2, mucus polymer concentrations drove complex viscosity (**Fig. 4A-C**) and showed a correlation with antibiotic treatment outcomes (**Fig. 4E and F**). Interestingly, when comparing the effects of HMW eDNA on the viscoelasticity of mucin solutions, we observed that at HMW eDNA concentrations ≥ 2 mg/mL, SCFM2 containing mucin had similar complex viscosities (**Fig. 4B-C)**. This result was similarly reiterated in the simulation model (**Fig. 4G**). This result indicates that eDNA dominates mucus viscoelasticity at disease-relevant HMW eDNA concentrations. In comparison, mucin had a lesser impact on viscoelasticity but was the major driver of antibiotic tolerance (**Fig. 4E and F**). When comparing the results shown in **Fig. 4A** to previous findings in both mucus harvested from human bronchial epithelial (HBE) cell cultures (47) and mucus recovered from endotracheal tubes (84), the data demonstrate that isolated commercial mucins do not fully capture the rheology of intact *ex vivo* mucus. *In vitro* systems are fundamentally unable to replicate all aspects of a host environment. Despite the fact that commercial mucins fail to capture the viscoelasticity of human respiratory mucus, our *in vitro* findings recapitulated the antibiotic tolerance of *ex vivo* treated sputum, suggesting that absolute viscoelasticity is not a dominant factor in defining variable antibiotic treatment efficacy across multiple subjects, although this was not directly tested. Despite the limitations of *in vitro* systems and commercial mucins, we demonstrated that an *in vitro* system, utilizing a modified version of SCFM2 with 2% w/v partially purified PGM and 1 mg/mL HMW eDNA, is a powerful tool for mimicking the necessary (though unknown) requirements to recapitulate antibiotic treatment outcomes in MAD *ex vivo* sputum.

The evidence that a single mucin concentration (2%) was able to recapitulate a majority of *ex vivo* tolerance was surprising given the heterogeneity in mucus concentrations observed among people with MADs (11, 42, 43). While 2% mucin was on average the best condition to capture the antibiotic tolerance in *ex vivo* sputum, other tested mucin concentrations also recapitulated antibiotic tolerance in some subjects. This is exemplified in the interactive simulation, which predicts that multiple combinations of mucin and HMW eDNA can achieve the same *P. aeruginosa* survival in response to antibiotic challenge.

Although the reasons for the failure of antibiotic therapy are multifactorial, controlling the environment by using the same *in vitro* growth conditions for all subject populations revealed a dominance of intra-subject *P. aeruginosa* diversity in driving antibiotic treatment failure across individuals with MADs. However, polymer-driven antibiotic tolerance is required to reveal the heterogeneity between subjects. SCFM2 containing 2% mucin and 1 mg/mL HMW eDNA was both necessary and sufficient to reveal the inter-population response of *P. aeruginosa* to tobramycin across subjects. Although mucin concentrations of < 2% significantly underestimated tolerance and > 2% significantly overrepresented tolerance across subject samples, the relative variability was inconsistent across other media conditions. The inter-population variability of *P. aeruginosa* in *ex vivo* antibiotic recalcitrance was best revealed in 2% mucin. (**Fig. S9B**). This indicates that while mucin and eDNA drive concentration-dependent effects on *P. aeruginosa* antibiotic recalcitrance, the effect is different between various isolates and populations of *P. aeruginosa*.

The physiological state of *P. aeruginosa* in the host respiratory mucus environment, combined with bacterial phenotypic and genetic diversification and adaptation within the airways, promotes tolerance to antimicrobial therapy (34–36). Although replicating the transcriptional signature of host-derived samples has been a primary focus of investigations into improving *in vitro* model systems, there has been limited examination of whether this translates to observed phenotypic outcomes, such as antibiotic recalcitrance (51, 53, 82). Refining well-defined *in vitro* systems beyond matching transcriptional signatures is necessary to accurately translate bacterial behaviors and phenotypes to their natural host environment.

Here we demonstrate that heterogeneity in what are known to be intra- and inter-diverse in host evolved populations drive differential responses to antibiotic challenge, phenotypes likely masked when looking at bulk population measurements, such as bulk transcriptomics. Single-cell traits, such as genetic and phenotypic diversity, transcriptional heterogeneity, and spatial organization (i.e., biofilms and aggregates), likely explain the proportion of the population that survives antibiotic treatment. Previous studies on the effects of mucin and eDNA concentrations on bacterial physiology have indicated that increasing mucin and eDNA concentrations increase the rigidity of biofilms (45) and promote the formation of multicellular aggregates (56, 57). The formation of concentration-dependent antibiotic tolerant biofilms and aggregates was not evaluated; however, previous studies identified polymer concentration thresholds for the formation of multicellular aggregates (56, 85–87). These findings may support the observation of a substantial shift in antibiotic tolerance between 1% and 2% mucin (**Fig. 2A and 5D**); this will be the focus of future studies.

The difficulty of finding participants to obtain fresh spontaneously expectorated sputum, along with the reduction in individuals producing spontaneously expectorated sputum, primarily due to the use of HEMT (88), increases the need for accurate *in vitro* model systems. A 2% mucin SCFM2 *in vitro* model provides an effective system to supplement or replace *ex vivo* sputum studies. A robust *in vitro* system that recapitulates the current standard for drug evaluation in *ex vivo* sputum has immense utility in translational investigative studies and novel therapeutic testing.

By exploring the effects of varying concentrations of the two dominant polymers in MAD mucus, mucin and eDNA, we were able to better understand how the host airway mucus environment impacts *P. aeruginosa* survival post-antibiotic challenge. Utilizing our knowledge of the concentration-dependent effects of mucin and eDNA on antibiotic tolerance of *P. aeruginosa*, we uncovered a unique *in vitro* condition capable of capturing the absolute value of tobramycin clearance efficacy, and, more importantly, the inter-subject variability of *P. aeruginosa* clearance in *ex vivo* treated sputum. We identified limitations of traditional laboratory growth medium (LB) and the original SCFM2 formulation (0.5% mucin) in recapitulating the inter-subject variability of *P. aeruginosa* sputum-derived clinical population response to antibiotic challenge. The information gained from this study begins to close the knowledge gap preventing successful treatment outcomes in chronic *P. aeruginosa* airway infections by understanding the contribution of the underlying lung microenvironment, in addition to bacterial physiology, in driving antibiotic treatment failure.

## Material and Methods

### Sputum Collection and Treatment

Expectorated sputum samples were collected from UNC Hospital’s Pulmonary Specialty Clinics. Sputum was collected with informed consent and UNC IRB approval (#02-0948). All participants were confirmed to be culture-positive for *P. aeruginosa* at a previous visit and were not prescribed Trikafta/Kaftrio modulatory therapy. Sputum was stored at 4°C immediately after collection and treated within 4 h of collection.

Prior to treatment, the sputum was homogenized and 50 μL of sputum was aliquoted into microfuge tubes. Samples were treated with 50 μL of vehicle control (PBS) or tobramycin (Sigma-Aldrich) at a final concentration of 300 μg/mL and gently tumbled at 37°C for 24 h. After 24 h, the samples were homogenized and treated with Sputolysin (dithiothreitol; Millipore Sigma) for 15 min at room temperature. Samples were pelleted, washed twice with sterile medium to remove trace antibiotics, serially diluted, and plated on *Pseudomonas* isolation agar (PIA; Thermo Scientific Remel). Agar plates were incubated at 37°C for up to 48 h to quantify the bacterial burden. Percent survival was calculated as the log-transformed ratio of tobramycin-treated to vehicle control-treated CFUs for each sample pair. *P. aeruginosa* grown on PIA from non-treated samples was collected, stocked in Lysogeny broth (LB) + 20% glycerol, and stored at −80°C as a total population present in the sputum.

### Antimicrobial Susceptibility Testing

Sputum population MIC values were determined using tobramycin Etest (Liofilchem) on cation-adjusted Mueller-Hinton agar plates (MHA; BD BBL) according to the manufacturers and Clinical Laboratory Standards Institute (CLSI) guidelines (60, 89). Briefly, MHA plates were spread with 1 × 10^8^ CFU/mL *P. aeruginosa* and a tobramycin Etest strip was sterilely added to the center of the plate. After 24 and 48 h of growth at 37°C, zones of inhibition were assessed. In the case of heteroresistance, the highest MIC was recorded. Clinical breakpoints for susceptibility and resistance adhered to the current CLSI guidelines (60), with the addition of a hyper-resistant breakpoint defined as MIC ≥ 10 times the tobramycin resistance breakpoint (8 μg/mL).

Broth MIC assays were performed using the microdilution method (89) in cation-adjusted Mueller-Hinton broth (MHB; BD BBL). Briefly, 5 × 10^5^ CFU were inoculated into 100 μL of media containing varying concentrations of tobramycin (Sigma-Aldrich), ciprofloxacin (Sigma-Aldrich), colistin sulfate (Sigma-Aldrich), and polymyxin B sulfate (Sigma-Aldrich). MIC assays containing mucin were performed using Muc5AC porcine gastric mucin (PGM; Sigma-Aldrich) dialyzed against cation-adjusted MHB. The MIC was visualized by the addition of 10 μL of 0.015% resazurin (Sigma-Aldrich) 24 h after incubation at 37°C and incubated for an additional 24 h (90).

### Bacterial Strains and Culture Conditions

*P. aeruginosa* reference strain, mPAO1 (91) was used in all *in vitro* experiments unless otherwise stated. *P. aeruginosa* was routinely cultured in LB or on LB agar (1.5% agar) and incubated under ambient air at 37°C. *P. aeruginosa* broth cultures were prepared from single colonies on a fresh agar plate and cultured in LB at 37°C with shaking at 250 rpm.

### Mucus Media Preparation

SCFM2 was prepared as previously described (49, 50) with some modifications. Briefly, a buffered base medium containing the same concentration of buffer, ions, and free amino acids used in SCFM was prepared, and the pH was adjusted to 6.8 with HCl (49). Muc5AC PGM (Sigma-Aldrich, cat # M1778) was used as the mucin source, dissolved in MilliQ water, and purified by ultracentrifugation (Beckman Optima LE-80L) via two 1 h spins at 48,000 × *g* to remove aggregated mucins and debris. After centrifugation, PGM was dialyzed against SCFM buffered base media using an 8 kDa molecular weight cut-off tubing to further remove impurities and equilibrate salt and metal concentrations. Mucin concentration after dialysis was quantified by measuring the total percent weight/volume (% w/v) solids and subtracting the added solids from the base media. The final components, including carbon sources, lipids, and DNA, were added last to complete the medium. Salmon sperm DNA (Sigma-Aldrich) was purified and sterilized using phenol:chloroform:iso-amyl alcohol (25:24:1) extraction. The completed medium was stirred at 37°C for at least 2 h to remove trace amounts of chloroform and tested for sterility prior to use. Reduced mucin was generated by treating purified PGM with 25 mM dithiothreitol (DTT; Sigma-Aldrich) and 4 M guanidine (Sigma-Aldrich; HMW, cat # D1626; LMW, cat #262012) for at least 3 h at 37°C in the dark. After 3 h, a fresh solution of iodoacetamide was added at 50 mM and incubated for an additional 30 min in the dark at 37°C to prevent the reformation of disulfide bonds. Reduced PGM was dialyzed against SCFM2 base to remove the DTT, guanidine, and iodoacetamide and completed as described above.

### Antibiotic Survival Assay

Prior to inoculation into SCFM2, overnight *P. aeruginosa* cultures were subcultured by dilution into fresh LB and cultured to an optical density at 600 nm (OD_600_) of 0.25. Mid-exponential phase *P. aeruginosa* cultures at an OD_600_ of 0.25 were inoculated into 100 μL media to a final inoculum of OD_600_ 0.0025 (1×10^6^ CFU/mL) in a 96-well plate. *P. aeruginosa* was incubated statically at 37°C under ambient air for 8 h. CFUs were quantified after 8 h of growth, and in a duplicate well, antibiotic was added and incubated statically at 37°C for an additional 24 h. Antibiotics were used at the following concentrations in all *in vitro* assays: tobramycin (300 μg/mL; Sigma-Aldrich), ciprofloxacin (300 μg/mL; Sigma-Aldrich), colistin sulfate (40 μg/mL; Sigma-Aldrich), and polymyxin B sulfate (40 μg/mL; Sigma-Aldrich). Antibiotic-treated wells were pelleted and washed twice with sterile medium prior to serial dilution to remove trace antibiotics. Percent survival was calculated as the log-transformed ratio of CFUs after antibiotic treatment to CFUs at the time of treatment (8 h). DNase experiments were performed with the addition of recombinant human DNase I (rhDNase I; Creative Biomart), at a final concentration of 3 μg/mL; the mean sputum concentration (92). The emergence of tobramycin-resistant isolates after antibiotic challenge was tested using a modified agar dilution method (89).

*In vitro* antibiotic survival assays with sputum populations were performed as described above with minor modifications. To maintain the population heterogeneity and avoid enrichment of isolates with the greatest fitness, sputum populations were passaged minimally and subcultured directly from frozen stocks. Agar plates of clinical isolates were incubated for an additional 24 h to count slow-growing isolates.

### Size-Exclusion Chromatography and Multi-Angle Laser Light Scattering (SEC-MALLS)

SCFM2 samples were diluted 1:250 in PBS, added to a chromatograph with a Superdex 1000 size exclusion column (15 × 2.5 cm), and eluted with 0.2 M NaCl and 10mM EDTA solution at a flow rate of 500 mL/min. The column effluent was passed through an in-line enhanced optimal system laser photometer (Dawn, Wyatt Technologies) coupled with a digital signal-processing interferometric refractometer (Wyatt/Optilab), which continuously measured the light scattering and sample concentration, respectively. The captured data were analyzed using Astra software (Wyatt Technologies).

### Antibiotic Diffusion

Texas Red-X succinimidyl ester (Invitrogen) was conjugated to tobramycin as previously described (93) and used to visualize diffusion. Fluorescently labeled tobramycin and similarly labeled dextrans (70 kDa) were added to the test media, mixed overnight, and inserted into capillary tubes preloaded with concentration–matched media. Diffusion of the conjugate through the capillary tube was measured every 15 min for 24 h using a Tecan Safire imaging plate reader. The diffusion coefficient for each conjugate through the test media was calculated from the intensity pattern, as follows:

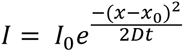

Where I_0_ is the background intensity, x is the direction down the tube, x_0_ is the position of the peak of the intensity pattern, t is the time at which the intensity pattern was imaged, and D is the diffusion coefficient.

### Particle Tracking Rheology

Viscoelasticity measurements were performed as described previously (7, 47, 84, 94). Briefly, 1 μm fluorescent microspheres with a COOH surface chemistry were added to the test media at a 1/500 – 1/1000 volume fraction and rotated overnight at 4°C. ∼ 5 μL of sample are then added to a sealed slide and imaged on a Nikon TE-U 2000 inverted microscope. 15-25 image sequences are collected using a Grasshopper 3 camera (FLIR) at 60 fps for 20 s. At least three slides were imaged per sample. Particle trajectories were tracked using TrackPy software, and the complex viscosity (*η*\*) was calculated from the Stokes-Einstein relationship using custom Python scripts, which are available upon request. The data were plotted as a swarm plot, with each dot representing the movement of a single particle.

### Modeling

We used JMP Pro (v17.2, SAS, Cary, NC, USA) to measure multivariate correlations, perform partial least squares regression, execute Monte Carlo simulations, and perform proportional analyses. Partial least squares regression used leave-one-out cross-validation to indicate a three-factor model. The X variables and interaction terms were included in the final model (all variable importance plot values were greater than 0.8). The prediction profiler and Monte Carlo simulations used mucin (% w/v) and DNA (mg/mL) inputs that reflected our experimental conditions: random values from a normal truncated distribution mean = 1.7, SD = 0.6, range = 0 – 5, and mean = 1.0, SD = 0.8, range = 0 – 4, respectively. Finally, we calculated the absolute least squared difference across all *in vitro* conditions versus the *ex vivo* condition per patient-derived culture for proportional comparisons. We defined differences from the *ex vivo* conditions (Y/N) as *p*-value ≤ 0.05 and used the resulting classification to perform analysis of means for proportions.

### Statistical analysis

All experiments were performed in at least biological triplicates across different media preparations. Statistical significance was assessed using Student’s two-tailed *t*-test, one-way analysis of variance (ANOVA), or analysis of means (ANOM), where appropriate. Statistical analyses were performed using GraphPad Prism version 10.1.0 for Windows (GraphPad Software, Boston, Massachusetts, USA). Differences were considered statistically significant at *p*-value ≤ 0.05.

## Data Availability

The interactive simulation model is publicly available at https://public.jmp.com/spaces/ALLUSERS/posts/3gyw4Hl2FCmqM648mJdGm/reports. All original data included in this study are available from the corresponding author upon reasonable request.

## Acknowledgments

The authors would like to thank the research participants and the members of the UNC CF Clinical Translational Core (Dr. Scott H. Donaldson, Margret Z. Powell, Dr. Kunal P. Patel, and Dr. Mary Leigh Anne Daniels) for providing and collecting sputum samples.

This research was supported by funding from the Cystic Fibrosis Foundation WOLFGA19G0 to M.C.W, HILL20Y2-OUT to D.B.H and the National Institutes of Health R21AI174088 to M.C.W, R01AG066710 and R01AG061188 to J.C.S. The UNC CF Clinical Translational Core and the Mucus/Mucin Biochemistry and Biophysics Core are funded by the National Institutes of Health NIDDK P30DK065988. M.A.G. was supported by funds from the National Health and Medical Research Council of Australia (NHMRC) through the NHMRC Synergy Funding Program (APP 1183640).

## Supplemental Figures

**Figure S1.**
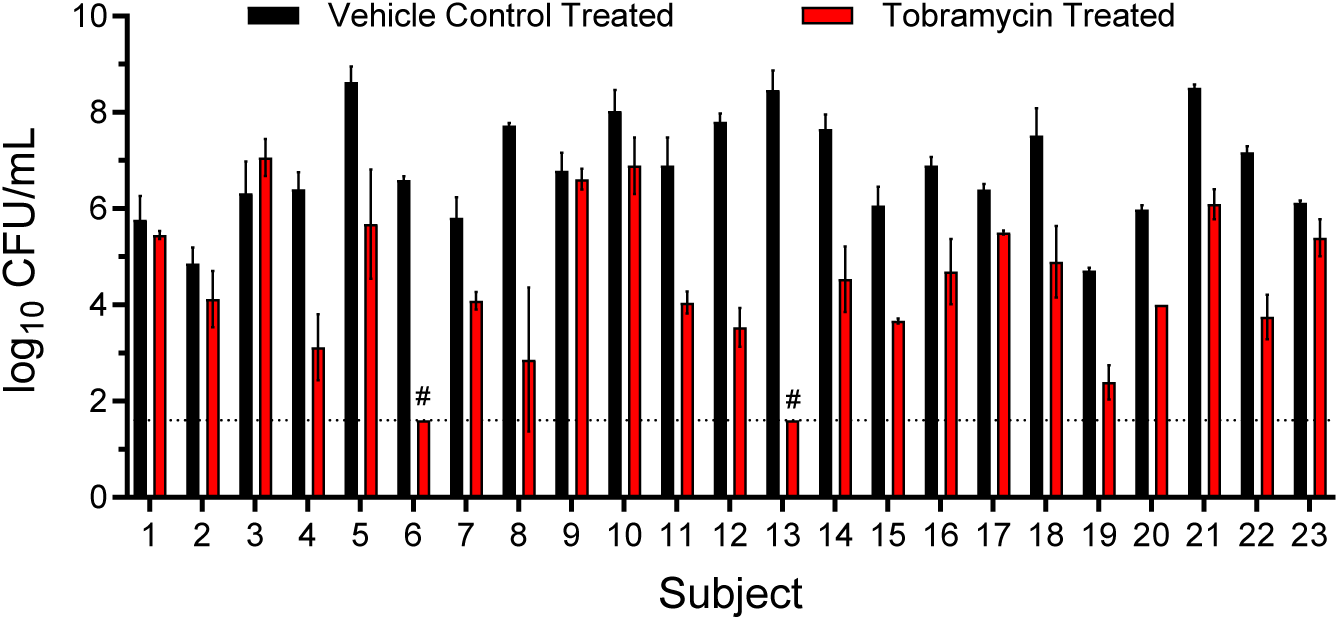
*Ex vivo* treated sputum CFU. CFU recovered on *Pseudomonas* isolation agar (PIA) from *n* ≥ 2 sputum aliquots (mean ± SD) after 24 h incubation with vehicle control (PBS) or tobramycin (300 μg/mL). # indicates no recovered *P. aeruginosa* after antibiotic treatment (limit of detection, 4 x 10^1^ CFU/mL).

**Figure S2.**
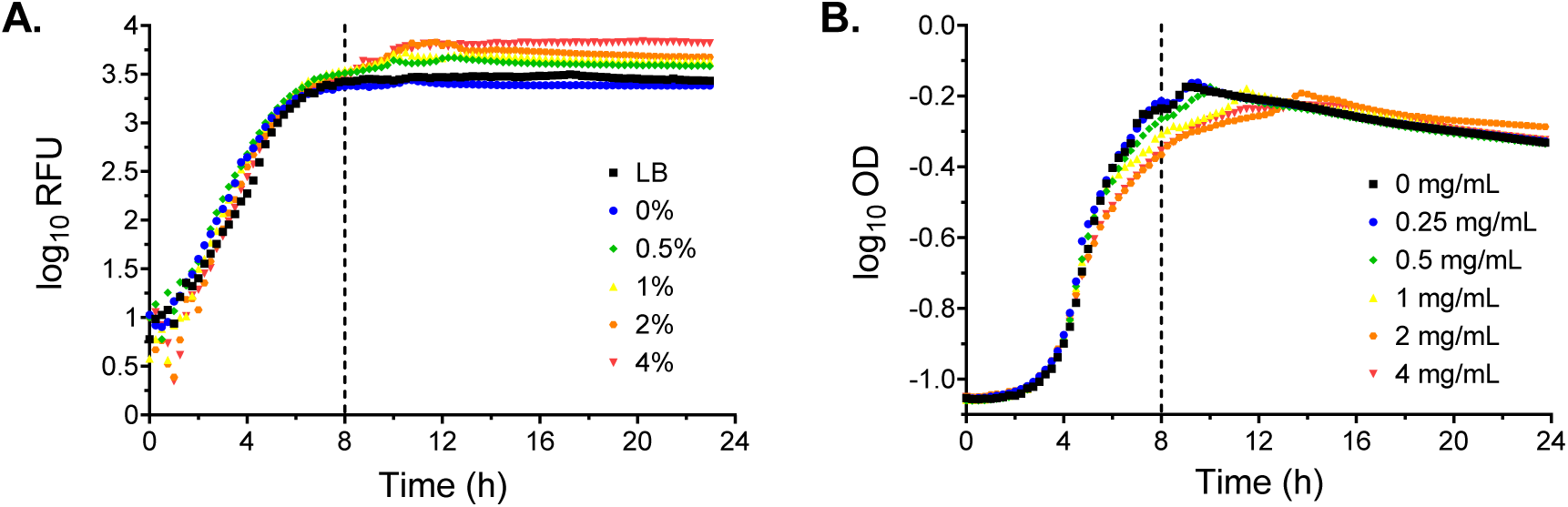
Mucin and eDNA do not affect growth kinetics of *P. aeruginosa*. **(A)** Growth curve of mPAO1 expressing *gfp*, in LB or SCFM2 with 0%, 0.5%, 1%, 2%, and 4% w/v mucin and no eDNA for 24 h. **(B)** WT mPAO1 growth curve in SCFM2 lacking mucin, with increasing concentrations of HMW eDNA. Fluorescence (483 / 535 nm) **(A)** or absorbance (600 nm) **(B)** measurements were taken every 15 min using a Tecan Infinite 200 PRO plate reader (Tecan Group Ltd., AG, Switzerland). Data are representative of the mean from *n* = 3 biological replicates in technical duplicate.

**Figure S3.**
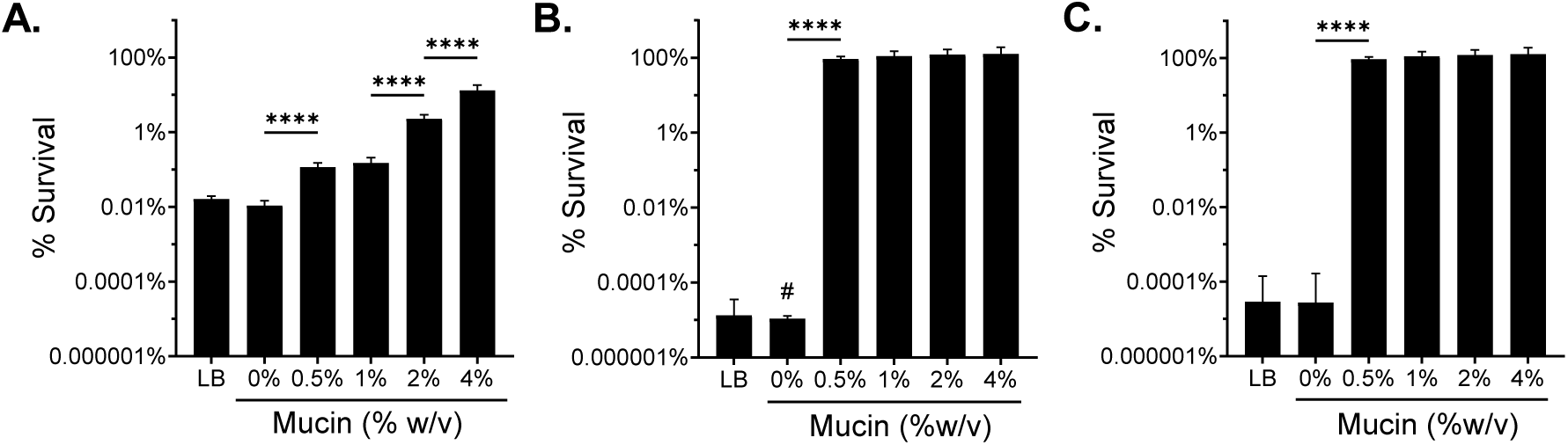
*P. aeruginosa* exhibits mucin concentration-dependent tolerance to multiple antibiotic classes. Survival of mPAO1 cultured in LB and SCFM2 media with increasing % w/v mucin and treated with **(A)** 300 μg/mL ciprofloxacin, **(B)** 40 μg/mL colistin, or **(C)** 40 μg/mL polymyxin B for 24 h. Mean ± SD of *n* ≥ 3 biological replicates. Statistical differences were analyzed using one-way ANOVA with Tukey’s multiple comparison correction. \*\*\*\**p*<0.0001. # indicates no recovered *P. aeruginosa* after antibiotic treatment (limit of detection, 10^2^ CFU/mL).

**Figure S4.**
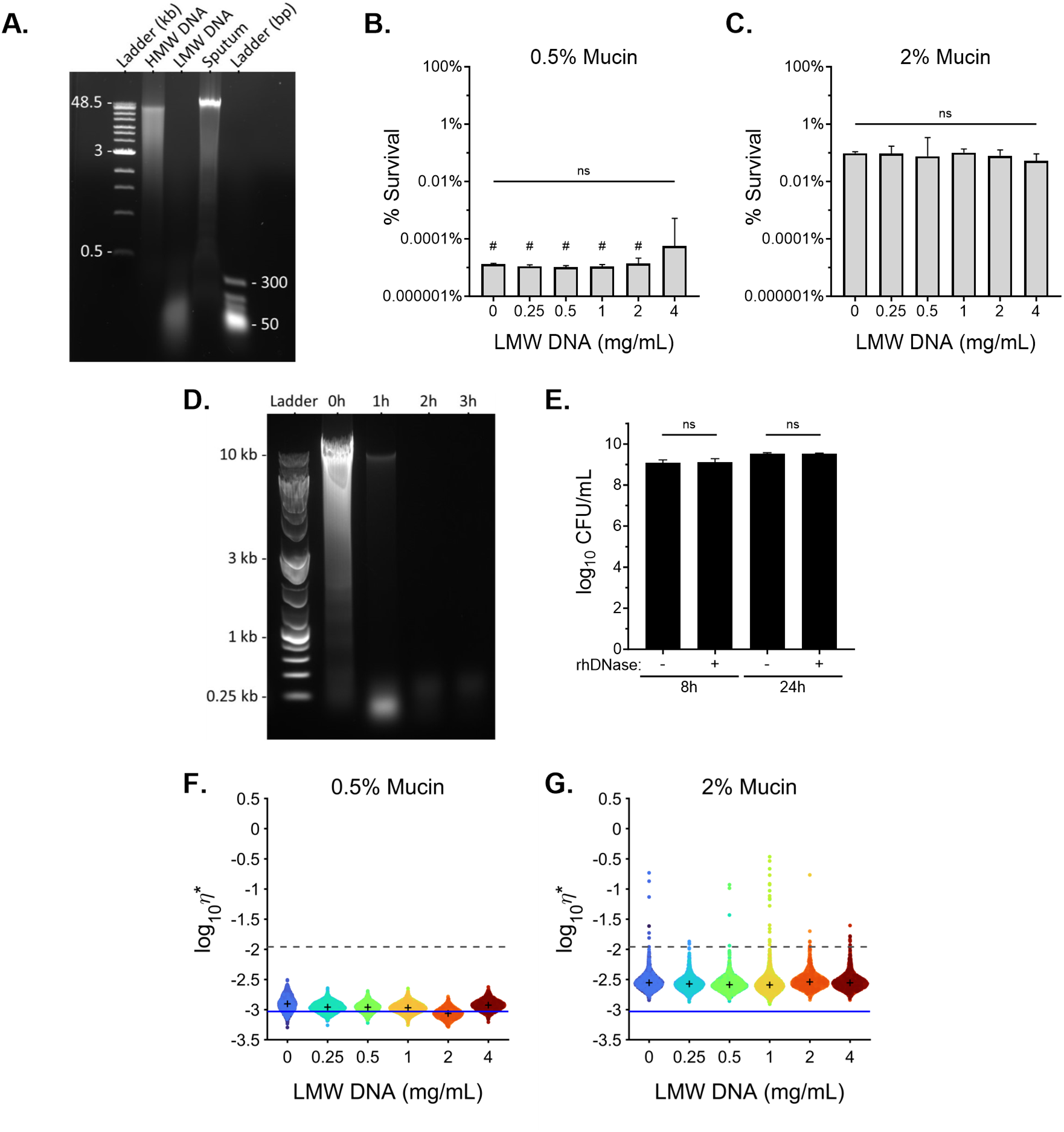
High molecular weight eDNA is necessary to impart tobramycin tolerance. **(A)** DNA gel electrophoresis of salmon sperm DNA from different commercial sources compared to a representative sputum from a CF subject. **(B-C)** % survival of mPAO1 after treatment with 300 μg/mL tobramycin in SCFM2 with **(B)** 0.5% or **(C)** 2% w/v mucin and increasing concentrations of LMW eDNA. Statistical differences were determined by one-way ANOVA. **(D)** Gel electrophoresis of SCFM2 containing 2% mucin and 1 mg/mL HMW DNA treated with rhDNase (3 μg/mL) for 0, 1, 2 or 3 h. **(E)** Effect of rhDNase (3 μg/mL) on the growth of mPAO1 in 2% mucin SCFM2 after 8 and 24 h. Mean ± SD of *n* ≥ 3 biological replicates. ns indicates not significant (*p*-value > 0.05). Probability distribution of each microbead *η*\* for increasing LMW eDNA concentrations in **(F)** 0.5% or **(G)** 2% w/v mucin SCFM2. Data are representative of *n* ≥ 3 technical replicates. The viscosity of water is indicated by a solid blue line and the complex viscosity at which mucus entangles and becomes gel-like (40) is indicated by a gray dashed line. + indicates the median complex viscosity for each sample.

**Figure S5.**
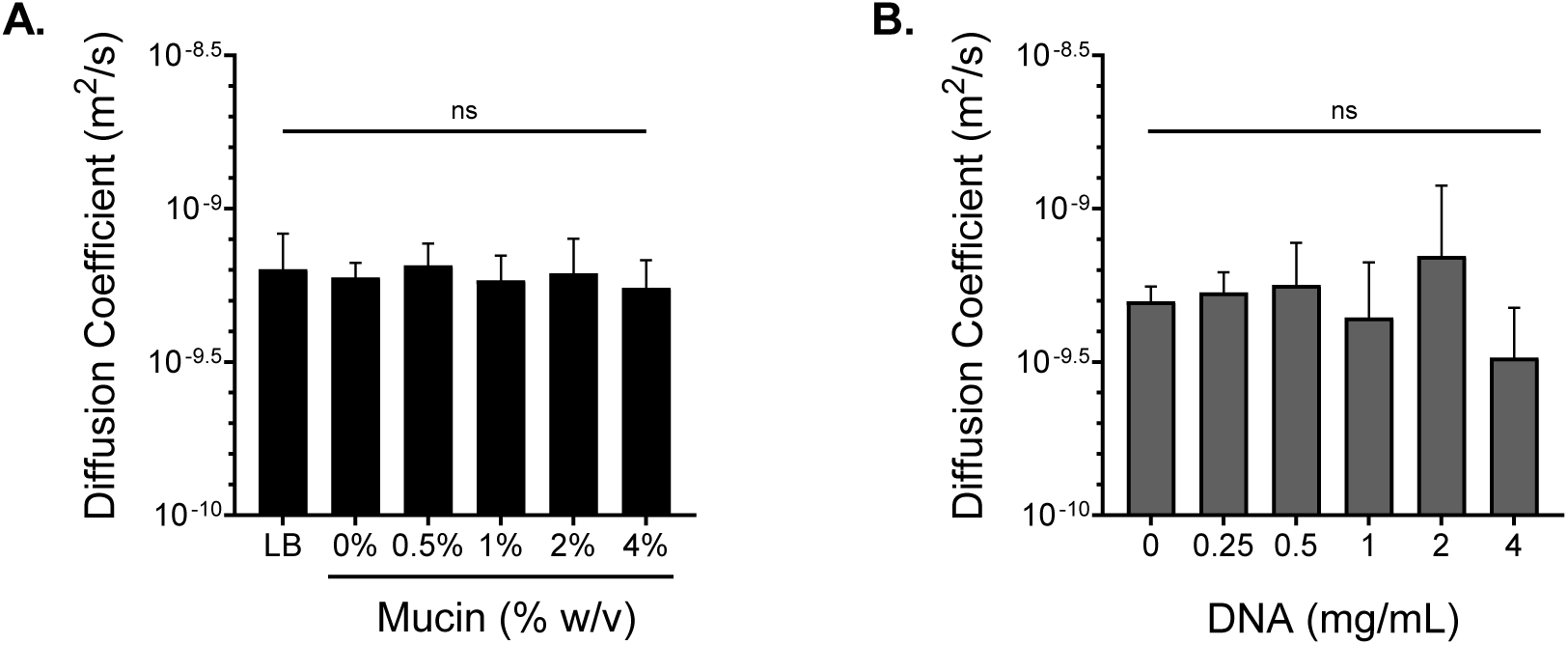
Antibiotic diffusion was not affected by polymer concentration. Diffusion rate of Texas Red conjugated tobramycin through **(A)** LB and SCFM2 with increasing mucin concentrations and **(B)** 0.5% w/v mucin SCFM2 with increasing concentrations of eDNA. Data are presented as the mean ± SD of *n* = 3 replicates. Statistical analysis was performed by one-way ANOVA. ns indicates not significant (*p*-value > 0.05).

**Figure S6.**
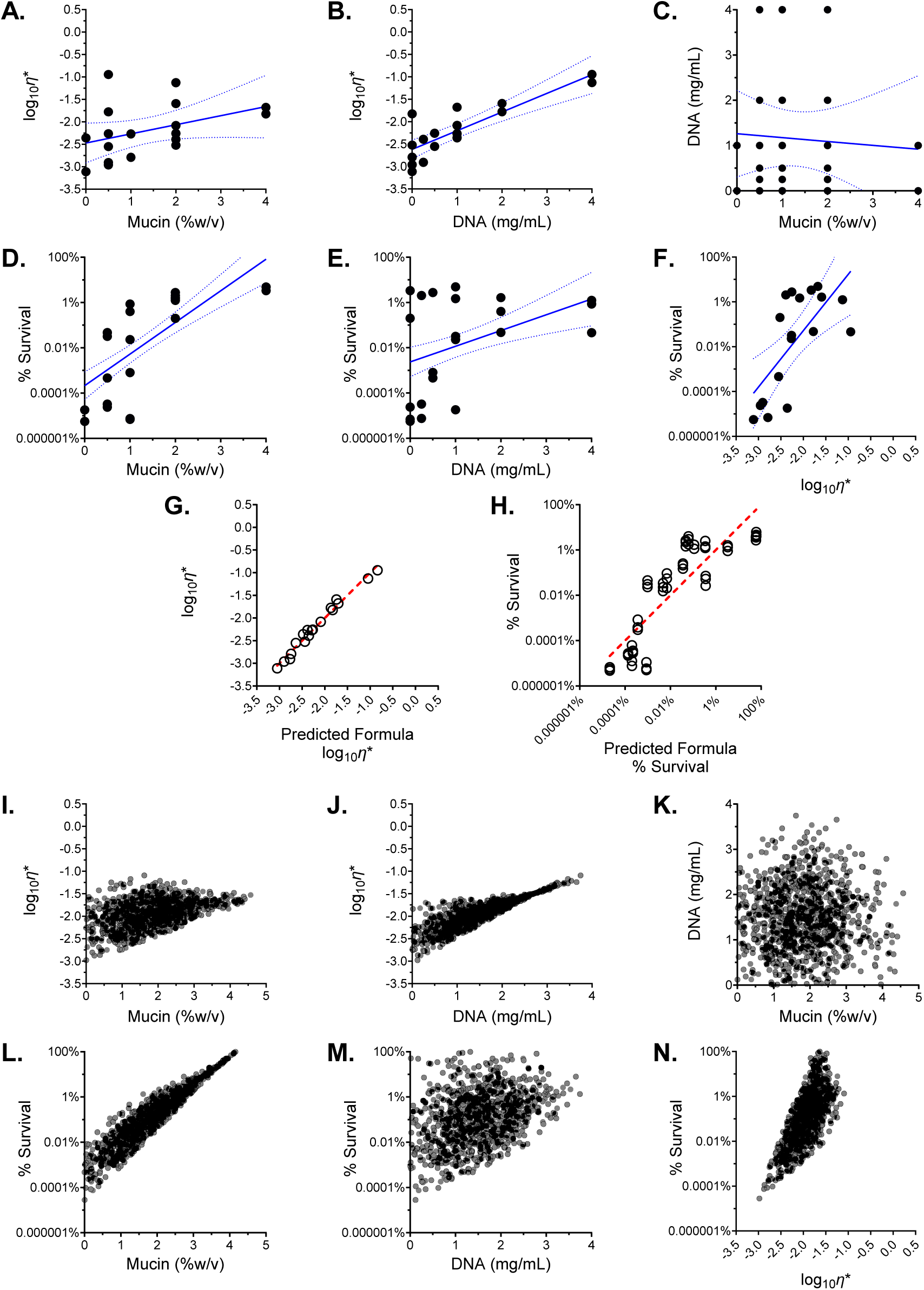
PLS regression model and simulation. **(A-F)** Linear trend and 95% confidence intervals of the relationship between the outcome variables (tobramycin survival and viscoelasticity) and input variables (mucin and eDNA concentration). **(G-H)** Three-factor model performance comparing the actual versus model predictions for **(G)** viscoelasticity and **(H)** antibiotic survival. **(I-N)** Individual biplot of 1000 simulated runs through the interactive model demonstrating the relationship between the outcome variables (tobramycin survival and viscoelasticity) and input variables (mucin and eDNA concentration).

**Figure S7.**
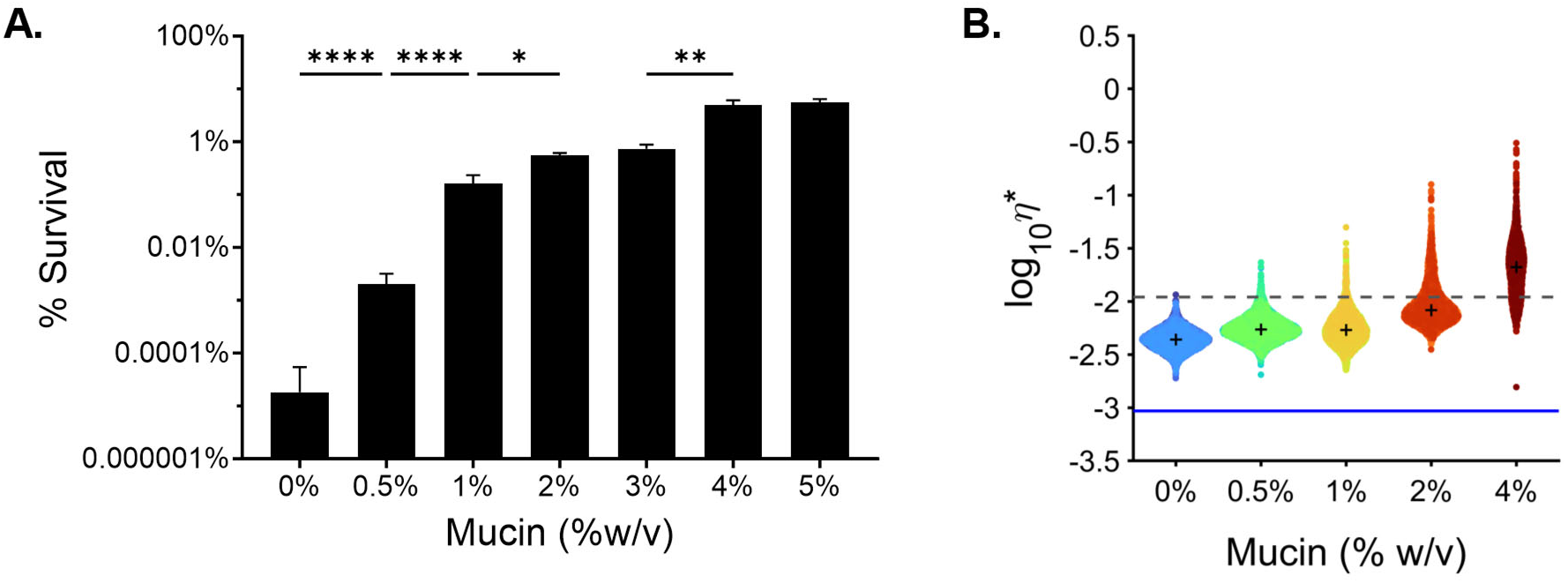
Antibiotic survival and complex viscosity of final media composition. **(A)** Tobramycin survival of mPAO1, and **(B)** complex viscosity of SCFM2 containing 1 mg/mL HMW eDNA and increasing % w/v mucin. Statistical differences were determined using one-way ANOVA with Tukey’s multiple comparison correction. \**p*<0.05, \*\**p*<0.01, \*\*\*\**p*<0.0001. Data represent the mean ± SD of *n* ≥ 3 biological replicates. The viscosity of water is indicated by the solid blue line, and the complex viscosity at which mucus entangles and becomes gel-like (40) is indicated by the dashed gray line. + indicates the median for each sample.

**Figure S8.**
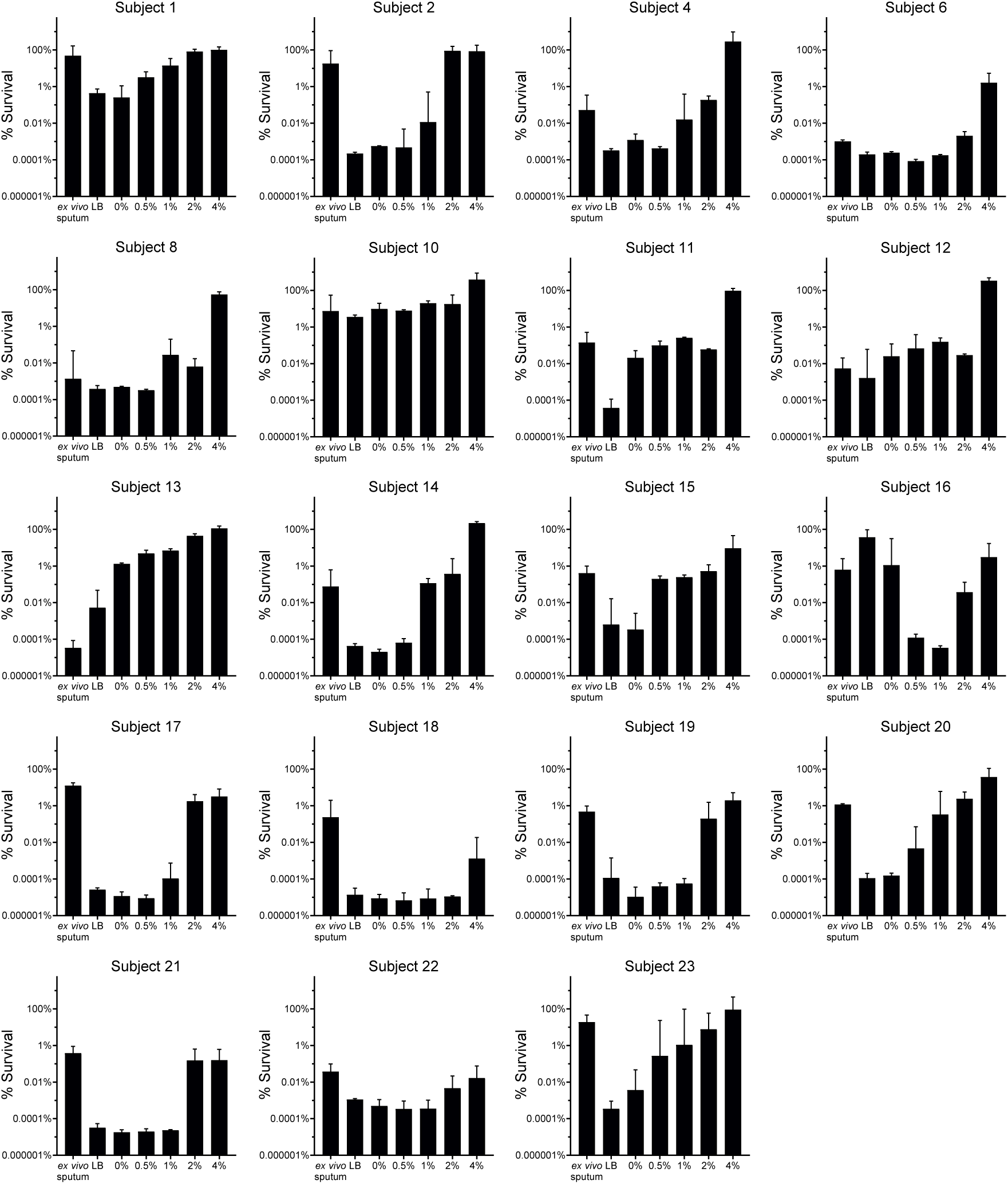
*In vitro* mucin concentration-dependent antibiotic survival of clinical populations. Percent survival of *ex vivo* treated sputum and *P. aeruginosa* populations recovered from each subject mock treated sputum evaluated *in vitro* for tobramycin tolerance in LB and SCFM2 containing 1 mg/mL HMW eDNA with increasing %w/v mucin. Hyperresistant sputum populations (MIC ≥ 80) were excluded. Data are representative of *n* = 3 independent replicates and plotted as mean ± SD. # indicates no recovered *P. aeruginosa* after tobramycin treatment (limit of detection, 10^2^ CFU/mL).

**Figure S9.**
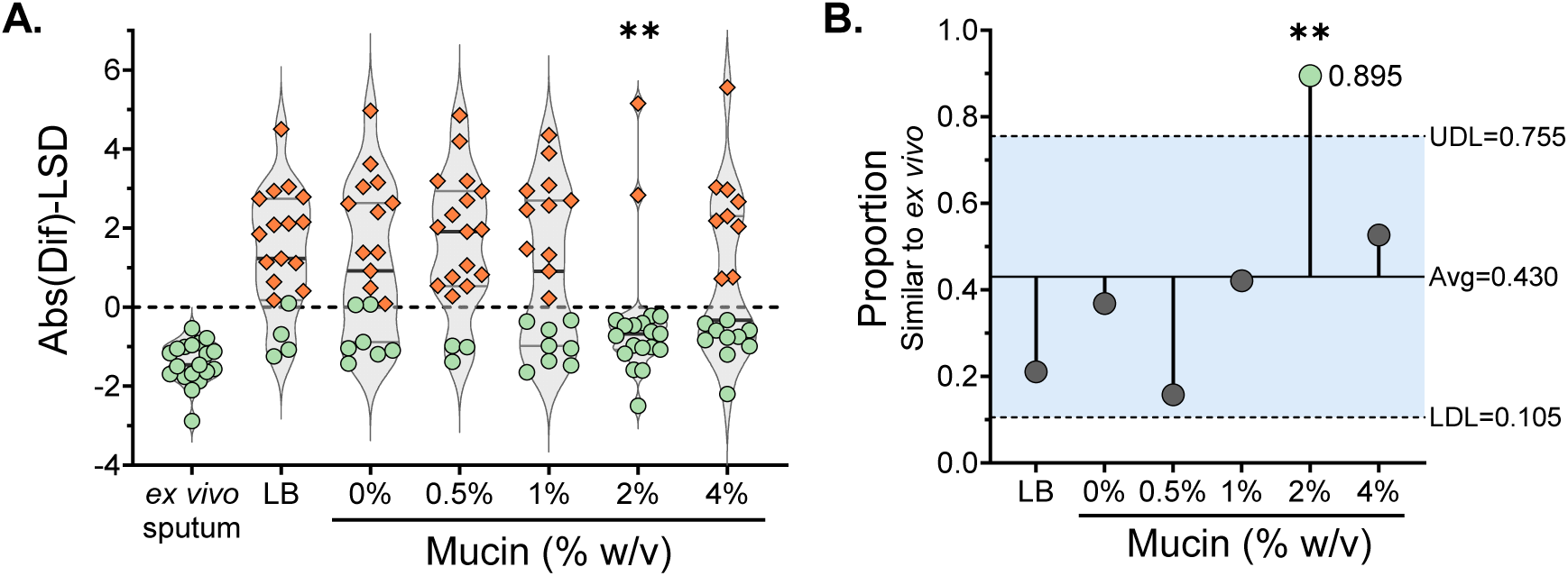
*In vitro* 2% mucin SCFM2 recapitulates *ex vivo* sputum antibiotic efficacy. **(A)** Comparison of the absolute difference between sample means minus Fisher’s least significant difference (Abs (Dif) - LSD). Values > 0 (orange circles) were considered different from *ex vivo* conditions at Dunnett’s post-test *p*-values < 0.05. **(B)** Proportion of *in vitro* cultures similar to *ex vivo* sputum tobramycin survival. Conditions that were disproportionate compared to the overall mean (43%) were identified via analysis of means for proportions (α = 0.01). The 2% mucin condition was the only condition above the upper decision line (UDL), indicating a difference from the mean across all conditions. \*\**p*<0.01.

